# Resolving phenotyping discordance with SPACEMAP, an integrated machine learning framework

**DOI:** 10.1101/2025.11.27.690631

**Authors:** Bassel Dawod, Arely Perez Rodriguez, Sebastian Diegeler, Eslam Elghonaimy, Megan Wachsman, Purva Gopal, David Hein, Paul H. Acosta, Andrew Jamieson, Gaudenz Danuser, Robert D. Timmerman, Satwik Rajaram, Todd A. Aguilera

## Abstract

Multiplex imaging technologies have revolutionized our ability to study cellular behavior within the tissue microenvironment. Translating this complex data into meaningful biological insights requires a unified analytical framework. To address this, we developed SPACEMAP (**S**patial **P**henotyping **A**nd **C**lassification with **E**nhanced **M**ultiplex **A**nalysis **P**ipeline), a comprehensive Python and Qupath-based platform for multiplex imaging analysis. SPACEMAP integrates image registration, segmentation, artifact removal, tissue and zone classification, spatial feature extraction, and a consolidated phenotyping approach into a single system. A core feature of SPACEMAP is its high-fidelity phenotyping. To evaluate classification performance, we benchmarked our method RESOLVE, against three established approaches, Leiden clustering, Self-Organizing Maps, and SCIMAP revealing substantial disagreement among them. SPACEMAP overcomes this through two complementary workflows: a machine learning model trained on expert-labeled cells, and a consensus classifier that integrates high-confidence cells across methods. Here, we validated SPACEMAP on in-house colorectal cancer samples and a public dataset, demonstrating its robustness.

## Introduction

Multiplex imaging technologies enable the simultaneous detection of dozens of markers in situ, providing a comprehensive view of cellular composition and spatial organization (1–4). There are multiple approaches but proteomic image-based technologies provide unique opportunities to incorporate subcellular positioning of proteins and the opportunity to extract multiple spatial features (5). Among these, Co-Detection by Indexing (CODEX) can visualize dozens of antigens within a single tissue section at single-cell and subcellular resolution through iterative cycles of ssDNA-barcoded antibody staining and fluorescent-labeled complementary ssDNA-reporter imaging (6). As these methods evolve, the scale and complexity of the resulting datasets create significant analytical hurdles, particularly in integrating molecular and histopathological information to ensure high-fidelity segmentation and robust, objective cell phenotyping (7–10). Extracting biologically relevant information from these data further requires multilayer image analysis pipelines, yet current tools often handle these steps in a disjointed manner, compromising reproducibility and analytical depth (11–13).

Within these pipelines, accurate segmentation of individual cells in densely crowded tissues remains a critical bottleneck (14). Pre-trained Deep learning-based tools that could be further trained for an application such as DeepCell (15) and Cellpose (16) address this, though their outputs still require validation. Additional pre-processing steps including image registration, artifact removal and data normalization are essential to preserve data fidelity and ensure comparability across samples (17). Finally, cell phenotyping, the assignment of cell identities based on marker expression, remains particularly challenging (18). Auto-fluorescent background in fluorescence-based multiplex imaging can interfere with accurate marker detection, further complicating phenotyping efforts. Moreover, algorithmic approaches often yield discordant classifications that can confound biological interpretations. Methods such as Leiden Clustering (19), Self-Organizing Maps (SOMs) (20) and SCIMAP (21) each offer distinct advantages but are limited by parameter sensitivity, biases in data assumptions and reliance on manual references inputs, ultimately limiting reproductivity and scalability.

To overcome limitations in the field such as fragmented image processing and discordant cell classifications that hinder harmonized analysis, we developed SPACEMAP. This comprehensive, modular computational pipeline unifies all critical steps of spatial analysis into a streamlined, scalable framework, consolidating foundational image processing steps, including co-registration hematoxylin & eosin (H&E) histology for anatomical context and single-cell segmentation. The core conceptual advance, however, is the multi-tool-phenotyping strategy, which leverages machine learning models trained on both manual reference cells and a novel reference-free consensus to deliver high accurate, reproducible cell classifications. By integrating these innovations, SPACEMAP establishes a new standard for extracting reliable biological findings from complex spatial proteomic data, advancing the study of the tumor microenvironment for both biological discovery and clinical translation.

## Results

### SPACEMAP facilitates image analysis into a unified workflow

To address the challenges of analyzing multiplexed imaging data, we developed SPACEMAP, a comprehensive and modular pipeline that integrates image registration, pre-processing steps, deep-learning segmentation, and robust phenotyping. We validated SPACEMAP on CODEX-stained colorectal cancer (CRC) tissue sections from patients treated on the INNATE trial who received neoadjuvant short course radiotherapy followed by chemotherapy and randomized to receive an anti-CD40 agonist with treatment.

The workflow encompasses key steps including CODEX-to-H&E image registration, cell segmentation, tissue annotation, artifact removal, batch correction, spatial feature extraction and phenotyping (Fig. 1a). We defined signatures to classify cells into 11 distinct types across non-immune, lymphoid, and myeloid lineages based on combinations of canonical markers (Fig. 1b). RESOLVE was developed to fully automate cell phenotyping using unsupervised marker threshold detection on individual marker intensity distributions to eliminate subjective steps. Cells were then assigned to one of 11 predefined types, with normalization and tie-breaking logic applied by assigning the phenotype to the highest probabilistic one (Fig. 1c). This automated, rule-based approach defines RESOLVE’s strength, enabling accurate identification of immune cells infiltrating in the dense epithelial layer (Supplementary Fig. 1). The outputs from RESOLVE and three established approaches Leiden clustering, Self-Organizing Maps (SOMs), and SCIMAP were then used to train two distinct machine learning workflows for final cell classification: a manual-reference model and a consensus model trained on cells consistently immunophenotyped by three of the four methods. Machine learning outputs were then evaluated using an independent reference dataset (Fig. 1d).

**Figure 1.**
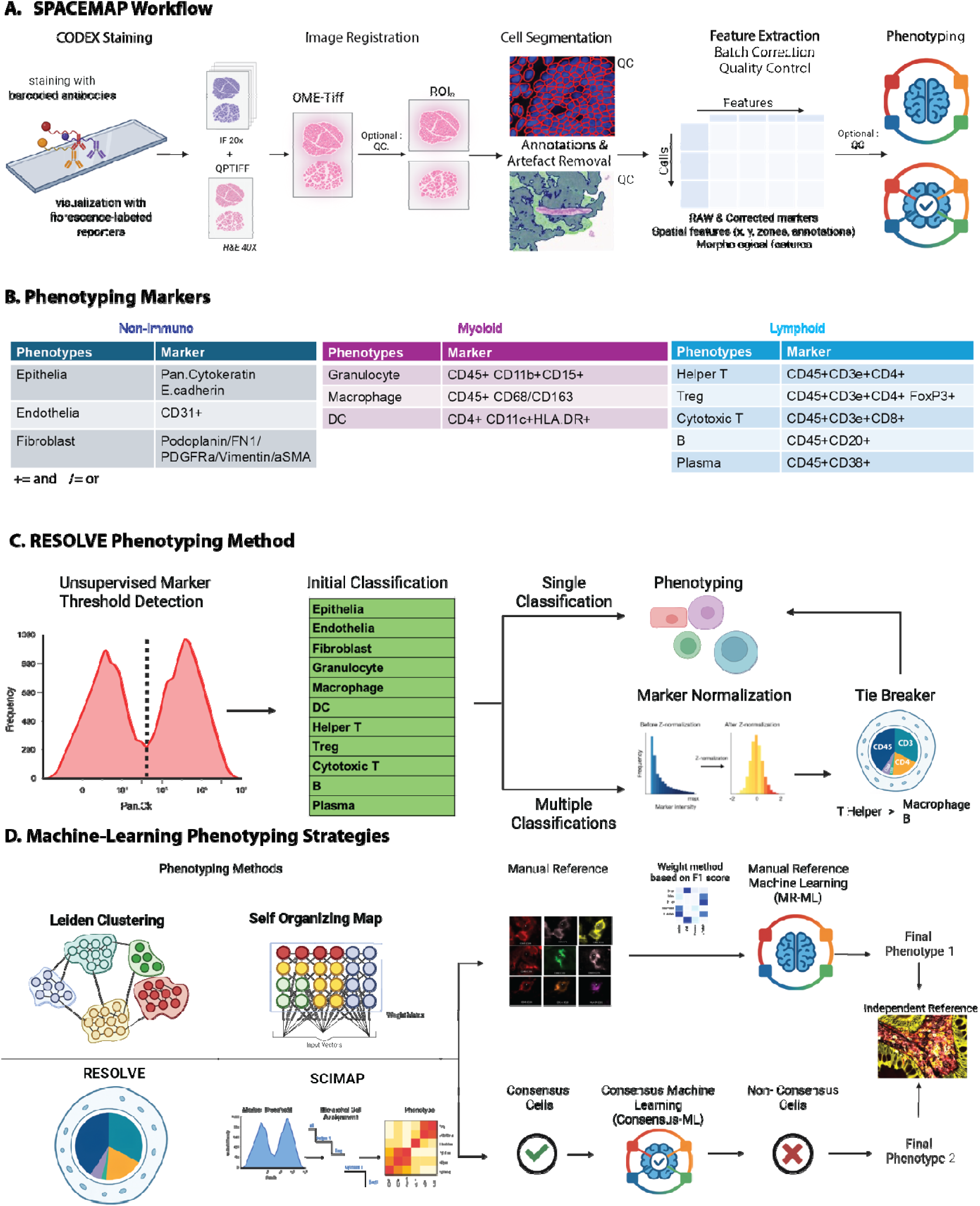
SPACEMAP integrates image registration, segmentation, annotation and phenotyping into a united workflow. **a** An illustration of the SPACEMAP analysis pipeline. **b** List of canonical markers used for phenotyping. **c** Overview of our new method RESOLVE: a fully objective phenotyping approach. **d** Schematic of two machine-learning driven phenotyping strategies in SPACEMAP using manual reference or consensus cells.

### Trained Segmentation enables personalization to tissue of interest

Image-based spatial biology workflows face several technical challenges, particularly in cell segmentation, where marker spillovers from neighboring cells and inaccurate detection of cell size can lead to erroneous cell phenotyping and inaccurate spatial analyses (9). Recognizing these limitations, SPACEMAP provides users the flexibility to select between two well-established deep learning-based segmentation models: DeepCell (Mesmer) (15) which employs convolutional neural networks trained on the TissueNet dataset for robust nuclear and cytoplasmic delineation in multi-channel tissue images, and Cellpose (16), a generalist algorithm utilizing gradient flow tracking and diffusion-based topological maps for versatile segmentation across diverse cell morphologies without requiring extensive retraining (Fig. 2a). Benchmarking studies indicate that Cellpose holds a slight advantage over DeepCell in overall precision and recall for fluorescent cell segmentation, particularly in nuclear detection across varying densities, though both methods exhibit trade-offs in cytoplasm versus nuclei accuracy (22). In our study, both default models exhibited over segmentation and false positives, often detecting cells without in-plane nuclei, especially in colonic epithelia. However, we observed that default DeepCell frequently over-segmented cells in regions with weak DAPI signal, while default Cellpose produced fewer artifacts (Fig. 2b).

**Figure 2.**
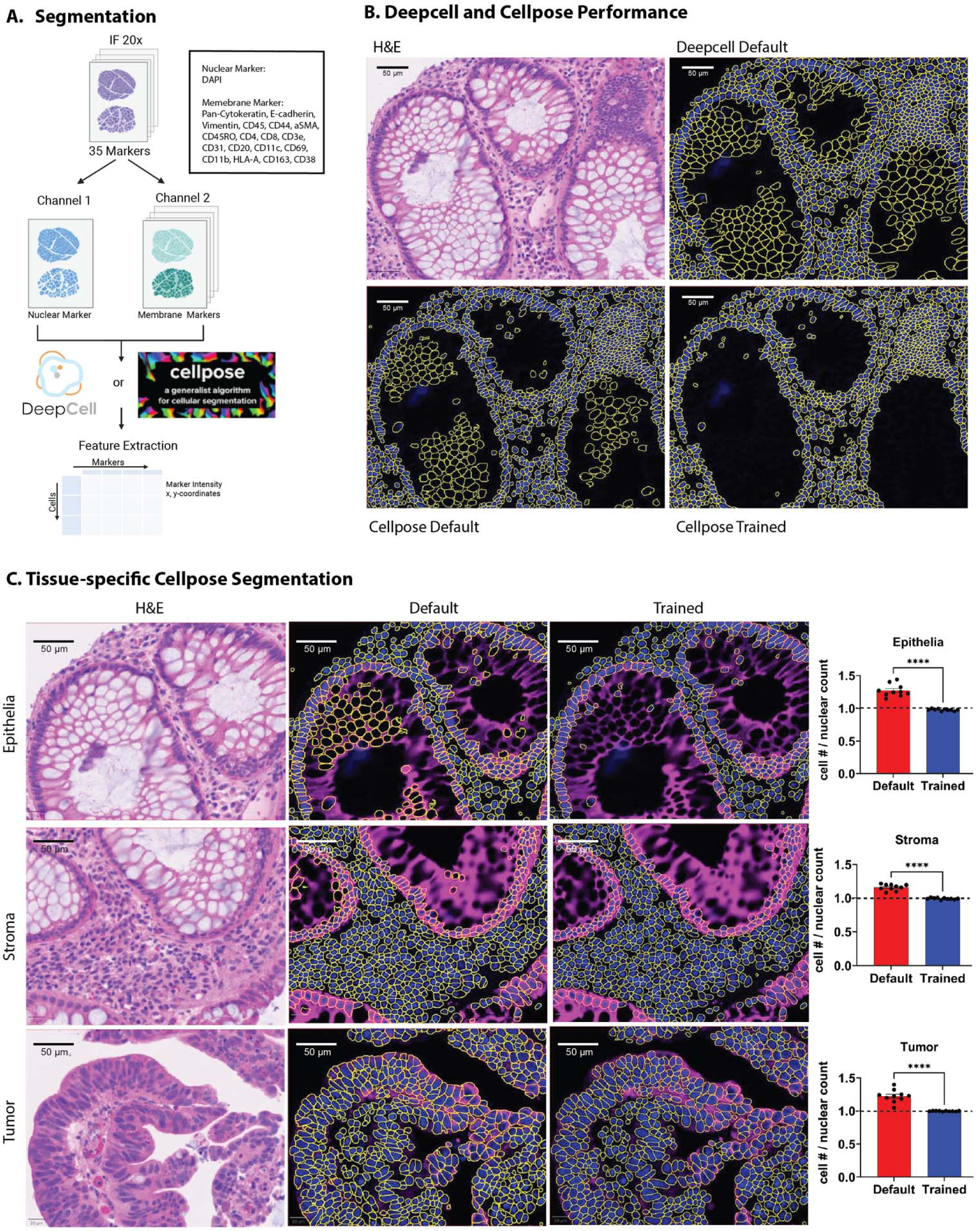
Cellpose with tissue specific training was selected over DeepCell for colorectal tissue segmentation. **a** Workflow for segmentation at 20x magnification using selected CODEX inputs, including a DAPI nuclear marker and a composite membrane channel (CD44, CD31, CD107a, E-cadherin, CD4, CD8, CD68, CD11c, aSMA, CD163). Segmentation can be performed via DeepCell (Mesmer) or Cellpose, followed by feature extraction of marker intensities, morphologies, and x-y coordinates. **b** Comparison of default DeepCell segmentation versus default Cellpose and trained Cellpose (yellow outlines on black background), along with H&E-stained tissue for reference. **c** Tissue-specific Cellpose segmentation performance in epithelial, stromal, and tumor regions, showing H&E references, default outputs, and trained outputs (yellow outlines on black background). Bar graphs quantify cell-to-nuclear count ratios (mean ± SEM; ***P < 0.001 by unpaired t-test), demonstrating significant reduction in over segmentation with the trained model (blue bars) compared to default (red bars). The black dashed line indicates the manual reference nuclear count used for comparison.

Given the observations detailed above, we pursued Cellpose for downstream analyses. To enhance accuracy, we trained a custom Cellpose model and benchmarked it against the default model using manual nuclear counts as reference. Across epithelial, stromal, and tumor regions, the trained model demonstrated significantly improved segmentation accuracy, with reduced over-segmentation and over-counting cells that more closely matched the nuclear counts than those generated by the default model (Fig. 2c). For scalability, the pipeline supports batch processing of multiple images and exports segmentation masks as QuPath-compatible GeoJSON files, a novel feature that facilitates crucial visual validation and supports the manual annotation of reference cells.

### SPACEMAP facilitates image processing and tissue annotations

Effective segmentation is a critical step towards robustly enumerating spatial data in complex environments, but the incorporation of multiple image processing steps is needed. Thus, SPACEMAP’s preprocessing pipeline incorporates image registration, ROI separation, background subtraction, pathologist-guided tissue feature classification, artifact removal followed by tissue mask generation, annotation-guided zone classification and feature extraction (Fig. 3a). In experiments combining H&E staining with CODEX multiplex imaging, SPACEMAP provides a novel registration tool to align and scale raw CODEX images with corresponding H&E sections, generating merged OME-TIFF files that integrate structural and molecular information for enhanced analysis (Fig. 3b). This multimodal fusion enables superior tissue annotation by leveraging complementary features, such as cellular morphology and architecture from H&E alongside multiplexed marker expression from CODEX, facilitating more accurate delineation of complex tissue compartments.

**Figure 3.**
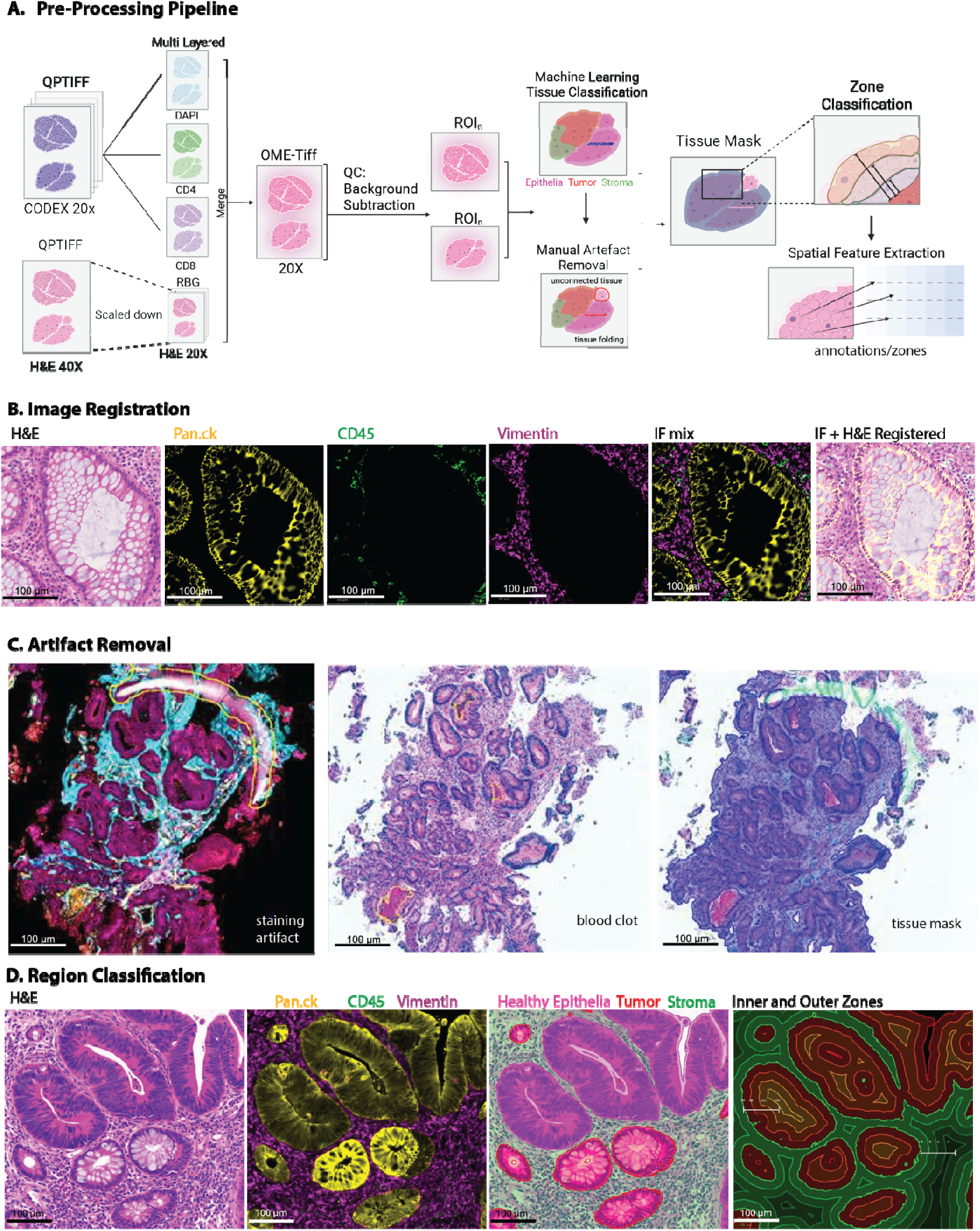
SPACEMAP’s preprocessing steps streamline data preparation for robust downstream analysis. **a** Schematic of the preprocessing workflow, from multi-layered QPTIFF inputs through merge, OME-TIFF generation, background subtraction, ROI delineation, machine learning-based tissue classification, manual artifact removal, tissue masking, zone classification, and spatial feature extraction. **b** Image registration examples, showing H&E (Pan.ck), CD45, Vimentin, IF mix, and IF + H&E registered overlays. **c** Artifact removal, with examples of staining artifacts, blood clots, and resulting tissue masks after exclusion. **d** Region classification on H&E, with overlays of Pan.ck/CD45/Vimentin, delineated healthy epithelium/tumor/stroma, and inner/outer zones.

For slides containing biopsies from multiple patients, SPACEMAP delineates distinct ROIs to split large images into separate files, reducing computational load, accelerating subsequent steps like artifact removal and annotation, while maintaining full compatibility with QuPath for visualization and the segmentation pipeline for feature extraction. Despite the built-in background subtraction, performed to correct for autofluorescence and non-specific signals, residual background may persist unevenly across markers due to imperfect removal. To mitigate this, SPACEMAP incorporates an optional yet highly recommended step using a dedicated blank imaging cycle acquired at the experiment’s end with non-specific fluorophores, enabling pixel-wise subtraction of lingering noise like uncorrected autofluorescence (Supplementary Fig. 2). Manual annotation then excludes artifacts such as staining errors, debris, blood clots, and dissociated tissue fragment unsuitable for spatial analysis (Fig. 3c).

Leveraging the registered H&E-CODEX data, a QuPath machine learning classifier, informed by pathologist expertise, partitions tissue into healthy epithelium, tumor, and stroma (Fig. 3d). Additionally, SPACEMAP generates spatial zones defining inner and outer regions based on proximity to annotations like tumors, enabling focused investigation of microenvironmental features (Fig. 3d). These preprocessing outputs, annotations and zones are seamlessly linked to cell-level features extracted via segmentation, such as marker intensities, morphology, and coordinates, enriching the dataset for downstream analyses including automated phenotyping and other potential spatial pattern quantifications. The complete workflow is illustrated for healthy and tumor tissues in Supplementary Figures 3 and 4 with all classification pathologist-validated against H&E images.

**Figure 4.**
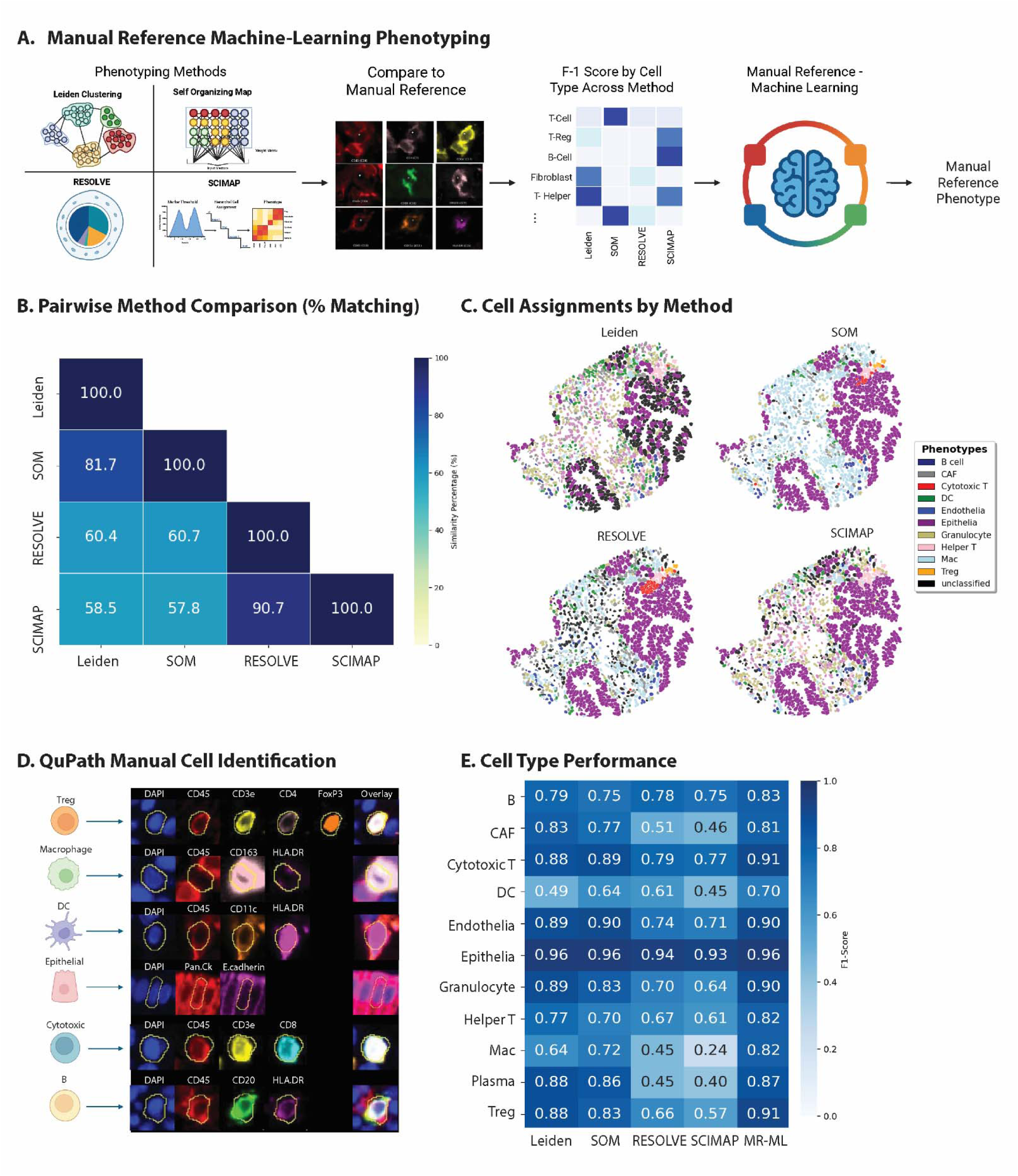
Benchmarking phenotyping approaches to inform a weighted manual reference-machine learning method. **a** Schematic of manual reference-machine learning (MR-ML) approach. **b** Pairwise comparison across methods: Leiden clustering, SOM, RESOLVE and SCIMAP. **c** Spatial plots of a representative region highlighting discordant cell identities across the four methods. **d** Representation of QuPath manual cell identification using canonical markers. **e** F1 score for each cell type across methods.

### Overcoming variability of immunophenotyping through machine learning

Following segmentation and feature extraction, we evaluated immunophenotyping methods to quantify variability in cell type assignments and developed an ensemble machine learning approach weighted by manual reference annotations (Fig. 4a). Pairwise comparisons across 2.6 million cells from the CRC biopsies revealed substantial discordance, with agreement ranging from 57.8% (SOM vs. SCIMAP) to 90.7% (SCIMAP vs. RESOLVE) (Fig. 4b). The elevated concordance between SCIMAP and RESOLVE likely stems from their shared threshold-based assignment frameworks, albeit with differing hierarchical phenotyping strategies. Spatial plots of a representative tissue region highlighted these assignment discrepancies (Fig. 4c). To resolve this variability and identify the most reliable classifications, we benchmarked Leiden clustering, SOM, RESOLVE, and SCIMAP against a manually annotated reference set generated via QuPath using canonical markers (Fig. 4d). F1 scores, metrics balancing precision and recall showed that all four methods accurately classified epithelial cells (>0.93), but performance varied widely for immune populations. For example, dendritic cell F1 scores were 0.49 for Leiden clustering and 0.45 for SCIMAP. For macrophages, RESOLVE scored 0.45 and SCIMAP scored 0.24 (Fig. 4e).

To mitigate these inconsistencies, we integrated outputs from all four methods into a manual reference-weighted machine learning (MR-ML) model, prioritizing contributions based on per-cell-type F1 score. This ensemble approach outperformed individual methods across all populations, elevating F1 score for challenging populations, such as dendritic cells (0.70) and macrophages (0.82), while maintaining high performance for abundant cell types like epithelial cells (0.96) (Fig. 4e). Additionally, Cohen’s Kappa, which is a robust statistic for assessing agreement unaffected by skewed class distribution, shows that manual reference- ML achieved the highest score, giving it approximately a 25% advantage over SCIMAP and a 10-20% advantage over other methods (Supplementary Fig. 5). Beyond this, the manual reference-ML method achieved overall highest scores for precision, recall and accuracy among all five evaluated methods (Supplementary Fig. 5).

### Consensus machine learning streamlines immunophenotyping

To overcome the time-intensive burden of manually annotating thousands of cells for a reference dataset, we developed a consensus-based workflow that integrates four complementary phenotyping algorithms. Cell phenotypes reached consensus if at least three methods agreed on their phenotype designation, representing 57% of our dataset.

A machine learning model trained exclusively on these consensus cells was then used to classify the remaining 43% of *“*non-consensus” cells. The model was trained on 75% of the consensus-cell dataset and validated on the remaining 25%, achieving a validation score of 0.98 (Fig. 5a). Performance of the consensus–machine learning model closely mirrored that of the consensus cells themselves. F1 scores showed the largest discrepancy in plasma B cells, while differences across other cell types were minimal (Fig. 5b). Spatial mapping further demonstrated strong concordance between phenotypes assigned by the manual reference–machine learning model and the consensus–machine learning model when applied to an independent reference tissue region not used in model training. Manual reference–machine learning achieved 85% accuracy against this independent reference, while consensus–machine learning achieved 84% (Fig. 5c). These values were comparable to the ∼85% agreement observed between the first manual reference used in the MR-ML training and the second manual reference employed for evaluation, reflecting normal variability between investigators (data not shown). The greatest divergence in F1 scores was observed in granulocytes (0.82 vs. 0.75), whereas all other cell types showed comparable performance (Fig. 5d).

**Figure 5.**
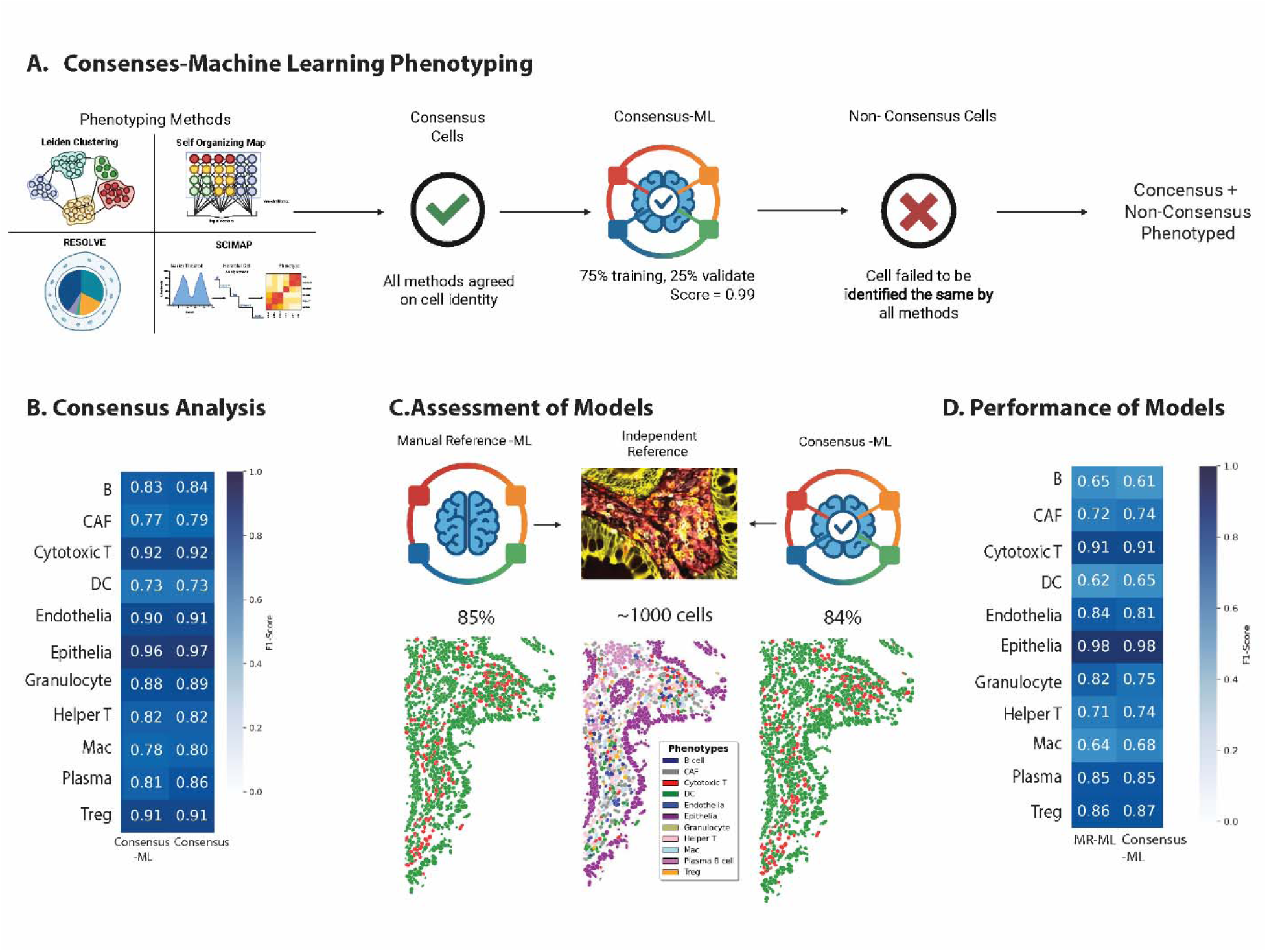
Consensus cell-based machine learning improves immunophenotyping efficiency. **a** Schematic of consensus-machine learning (consensus-ML) approach. **b** Comparison of F1 scores by cell type for consensus cells and consensus-ML. **c** Evaluation of both models on an independent reference tissue region. **d** Comparison of F1 scores by cell type for MR-ML versus consensus-ML on the independent tissue region.

### Reproducibility of the consensus–machine learning approach

To test reproducibility, we applied SPACEMAP’s consensus–machine learning workflow to an independent CODEX dataset published by Schürch et al. (6). We classified their cell phenotypes into nine distinct identities using canonical markers and manually phenotyped ∼ 2,500 cells to serve as a reference standard. After processing with our consensus–machine learning algorithm, 88% of cells matched the manual reference phenotypes (Fig. 6a). In contrast, the four individual methods Leiden, SOM, RESOLVE, and SCIMAP achieved lower concordance with the manual reference at 69%, 69%, 71%, and 81%, respectively (Fig. 6b). Pairwise comparisons revealed only moderate agreement between individual methods, with the highest concordance observed between SOM and Leiden at 79.7% (Fig. 6c). Comparing these methods on the phenotype level, best overall performance was observed in the consensus-ML method, showing balanced F1 scores across all phenotypes, with tumor, cytotoxic T cells and endothelial cells reaching 0.95, 0.86 and 0.89. Notably, B cells and granulocytes show high concordance with consensus-ML (0.73 and 0.80, respectively), while being under-recognized by RESOLVE and SCIMAP (B cells: 0.35 and 0.25, respectively), and SOM and RESOLVE (0.49 and 0.44, respectively (Fig. 6d). It is important to note that although SCIMAP yielded the highest matching percent (81%) to the manual reference, this was primary driven by its performance in tumor cell classification, as tumor cells compromised ∼50% of the dataset. Thus, the elevated accuracy did not clearly reflect superior performance across all phenotypes and shows that cell level precision needs to be evaluated as well. Comparing all phenotyping methods using multi-parameter performance evaluation, including recall, precision, accuracy, F1 score and Cohen’s kappa shows that consensus ML out-performs all other methods (Supplementary Fig 6). Notably is an advantage of consensus-ML in Cohen’s kappa of approximately 25% over SOM and RESOLVE and a 23-15% advantage over Leiden and SCIMAP respectively (Supplementary Fig 6).

**Figure 6.**
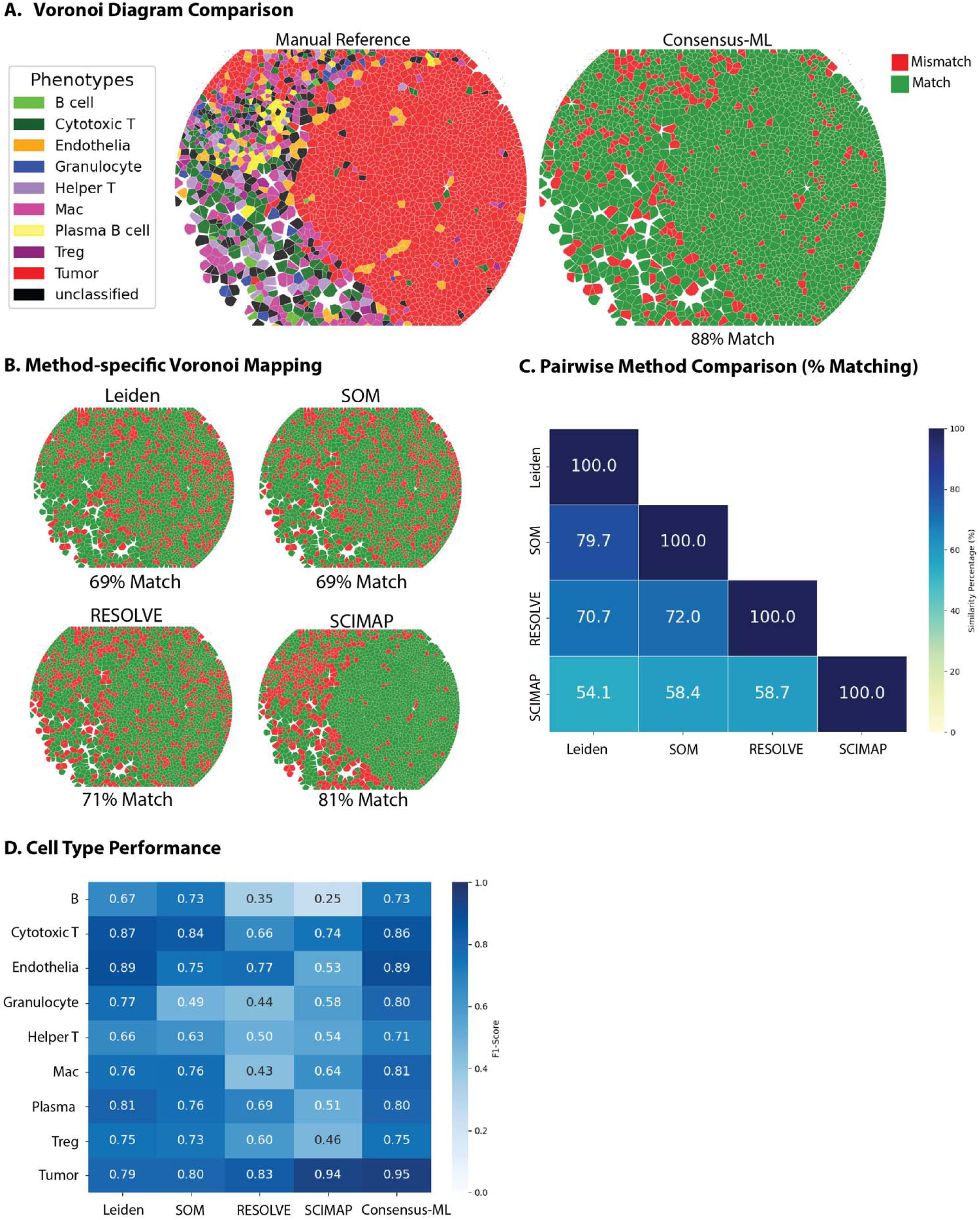
Public datasets confirmed the robustness of our consensus-machine learning approach. **a** Voronoi Diagrams highlighting agreement and disagreement between manual reference and consensus-machine learning. **b** Voronoi Diagrams illustrating method-specific discrepancies across Leiden, SOM, RESOLVE and SCIMAP. **c** Pairwise comparison across methods: Leiden clustering, SOM, RESOLVE and SCIMAP. **d** Comparison of F1 scores by cell type across individual methods and consensus-ML.

## Discussion

In this study, we introduce SPACEMAP, a comprehensive computational pipeline designed to harmonize and standardize multiplexed tissue imaging data. Our investigation reveals that rigorous computational analysis of such data necessitates adaptive workflows to mitigate variability on phenotyping outcomes. Through iterative refinement, we discovered that combining individual cell phenotyping methods with machine learning approaches can reconcile disparate results, yielding robust and accurate cell classifications in an unbiased manner. Much work has been focused on developing supervised (SCIMAP) and non-supervised (Leiden, SOM) approaches with some focusing on expert subjective calls (6) Even others have used a mix of cell segmentation and pixel-based classification to deal with the dense stromal heterogeneity (23). Altogether, we observed that a profound challenge persists in spatial biology: the absence of a universally accepted method for cell phenotyping has led to “method-dependent” variability in classification outcomes. This is compounded by the difficulty making workflows streamlined and removing as much risk of subjective bias as possible. Our findings confirm such limitations, highlighting how distinct algorithms can produce vastly different classifications on the same dataset. Notably, while methods excel with abundant and phenotypically distinct populations like epithelial cells, they falter with immune cell subsets, underscoring the need for a more comprehensive approach to address these discrepancies. By acknowledging the pitfalls of relying on a single algorithm, we underscore the imperative of developing alternative methods that can effectively tackle complex tissues such as the tumor microenvironment.

To propose a robust solution, our weighted model leverages the collective strengths of multiple algorithms. By learning which method is most reliable for each specific cell type, the model combines their outputs into a single, more accurate reference-free classification. The meaningful improvement in F1 scores demonstrates that this integrated approach is superior to any single method alone. This provides a reliable foundation for downstream spatial analyses. Although accurate, our first model trained on a manually curated reference set relied on a time-consuming and labor-intensive process. This led us to propose the consensus-based training strategy. By using agreement among multiple algorithms as a reference, we eliminate the need for a manual reference set. The high concordance between the consensus-trained model and the manual reference-machine learning model validates this approach with in-house and a public data set. This finding is significant because it provides a practical, scalable, and automated workflow for achieving high-fidelity immunophenotyping in large datasets.

Furthermore, our work underscores the importance of a rigorous, end-to-end workflow. Accurate cell phenotyping is contingent upon high-quality upstream processing. Through a combination of H&E registration, robust artifact removal, background subtraction, and optimized cell segmentation, SPACEMAP establishes a solid foundation for reliable downstream classification. It offers a validated, and scalable solution for analyzing complex spatial proteomics data and its weighted and consensus-based machine learning strategies directly address the critical challenge of method-dependent variability, providing more reproducible and reliable insights in spatial biology.

Despite these advantages, SPACEMAP has several limitations. In densely packed regions or at the edges of imaging planes, cells may be misclassified or missed. The consensus-based approach assumes that agreement among multiple algorithms reflects the correct cell identity, which may underrepresent rare or atypical phenotypes. Segmentation performance also varied in colorectal tissue, reflecting the complexity of epithelial architecture and stromal density. Additionally, dedicated spillover-correction tools such as REDSEA have been developed to address marker contamination between adjacent cells (9). Although earlier versions were technically challenging to implement reliably in dense tissues, ongoing improvements in these algorithms represent an important future direction. Integrating such tools into SPACEMAP could further enhance segmentation accuracy, particularly in highly compact or heterogenous regions. Because we select cells containing nuclei, standard histologic sections can include fragmented cells, particularly within colorectal epithelium that are not accounted. Future 3D imaging approaches will be critical to capture the true spatial architecture of tissues, enable more accurate quantification of cell–cell interactions, reveal migration dynamics, and refine morphological analyses. Finally, although SPACEMAP provides a flexible framework for spatial data integration, expert validation remains essential. Pathologist input will be important for diverse tissue types to ensure biologic relevance, and technical expertise will be required when extending the method to new imaging platforms. Nevertheless, SPACEMAP offers a generalizable framework that can guide future spatial mapping efforts across tissues and disease contexts.

Looking ahead, SPACEMAP is positioned to facilitate future research in spatial biology by enabling reproducible, large-scale studies across diverse datasets and platforms. By reducing reliance on labor-intensive manual annotations and standardizing workflows, SPACEMAP makes high-fidelity phenotyping scalable and accessible. Its modular design allows adaptation to emerging imaging technologies and marker panels, while reducing technical variability and allowing researchers to focus on biological discovery, accelerating efforts to map tissue microenvironments and guide therapeutic strategies.

## Methods

### Datasets

The primary dataset for this study was derived from the INNATE trial (NCT04130854), a Phase II randomized clinical trial for patients with Stage II/III locally advanced rectal adenocarcinoma. Tumor tissue samples were collected at pre-treatment and post-Short-Course Radiotherapy (SCRT) with or without the anti-CD40 agonist sotigalimab. These samples were the baseline input for primary analysis. To validate the robustness, generalizability, and reproducibility of our custom analytical pipeline, SPACEMAP, we applied it to the publicly available single-cell spatial proteomics dataset from Schürch et al. (“Coordinated Cellular Neighborhoods Orchestrate Antitumoral Immunity at the Colorectal Cancer Invasive Front”)(6). This external dataset, based on Tissue Microarray (TMA) of colorectal cancer was used to verify the phenotyping accuracy and the overall functional integrity of SPACEMAP, thereby increasing confidence in its capability to achieve consistent results and maintain generalizability for future spatial analyses.

### Antibody Validation

All antibodies were validated before use on INNATE samples using two FFPE multi-tissue microarrays (TMAs) containing 2-mm cores from 16 tissue types: six normal tissues (tonsil, lymph node, spleen, pancreas, colorectal), colorectal carcinoma (CRC), high-grade pancreatic intraepithelial neoplasia (PanIN), six pancreatic ductal adenocarcinomas (PDAC), and three matched PDAC lymph-node metastases. Tumor and normal regions were annotated on corresponding H&E-stained sections by a board-certified pathologist (M.B.W.).

Antibody validation was based on four predefined criteria: (1) correct subcellular localization, (2) co-staining with known markers, (3) appropriate counter-staining, and (4) expected tissue-specific staining patterns. Expected localization for each marker was determined from the literature (e.g., nuclear Ki67; membrane CD8). Each validation round included markers co-expressed with the target marker (e.g., CD3 with CD8 on cytotoxic T cells) and markers that should be mutually exclusive (e.g., CD31 absent on CD8^⁺^ cells). When needed, tissue-specific expression was confirmed using representative tissues—such as high Ki67 expression in CRC epithelial glands co-staining with PanCK and excluding CD31^⁺^ vasculature. Observed staining patterns were compared with established IHC reference images form proteinatlas.org or literature to ensure concordance.

Once all criteria were met, staining conditions for INNATE samples were optimized to improve signal-to-noise and refine antibody dilutions. A final verification step on optimized INNATE sections confirmed expected co- and counter-staining patterns. Necrotic regions, which can exhibit nonspecific reporter fluorescence independent of antibody binding, were identified by unstructured high-intensity signal and confirmed on H&E by a pathologist. These regions were excluded from analysis.

### Antibody conjugation

Unconjugated antibodies (Table 1) were barcoded using Akoya oligonucleotide conjugation kits (Cat# 7000009). Briefly, 50 µg of purified antibody was concentrated on a 50 kDa centrifugal filter (Amicon Ultra 0.5, Cat# UFC505024, Sigma Aldrich) and reduced for 30 min at room temperature using the provided Reduction Mix. After centrifugation (12,000 g, 8 min) to remove the reduction buffer, the filter was equilibrated with Conjugation Solution and lyophilized barcodes were reconstituted and incubated with the reduced antibody for 2 h at room temperature. Excess barcode was removed by three washes with Purification Solution (12,000 g, 8 min each). Conjugated antibodies were eluted in Antibody Storage Solution by inverting the filter and centrifuging at 3,000 g for 2 min.

### Multiplex CODEX Immunofluorescence Labeling

FFPE tissue sections (5 µm) on Superfrost Plus slides were baked at 60 °C overnight, deparaffinized in HistoChoice (2×5 min), cleared in 100% ethanol (5 min), rehydrated through graded ethanol (90–30%; 5 min each), and rinsed in ddH□O (2×5 min). Antigen retrieval was performed in 1× AR9 buffer (pH 9) under high pressure for 20 min, followed by cooling to room temperature and ddH□O washes. Autofluorescence was quenched by two 45-min incubations in photobleaching solution (4.5% H□O□, 20 mM NaOH in PBS), followed by PBS (4×5 min) and hydration buffer washes.

Slides were equilibrated in staining buffer for 20 min, then blocked with staining buffer containing 4.75% (v/v) Blocker N, J, S, and G2. Antibody cocktails (200 µl/slide; Table 1) were applied within a hydrophobic barrier and incubated for 3 h at room temperature in a humidity chamber. After staining, slides were washed in staining buffer (2×2 min) and fixed in post-stain fixation solution (1.6% v/v in storage buffer) for 10 min, rinsed in PBS, and fixed sequentially in cold methanol (−20 °C, 5 min) and final fixation solution (Akoya Fixative Reagent 1:50 in PBS, 20 min). Slides were rinsed in PBS and stored in storage buffer until Flow Cell assembly.

INNATE samples were stained with 37 antibodies, of which 34 passed QC and were used for downstream analysis.

### CODEX Acquisition

Images were acquired on a PhenoCycler-Fusion 1.0 (Akoya Biosciences) following manufacturer instructions. Reporter plates were prepared such that each well corresponded to one imaging cycle; wells for the first and last cycle contained reporter buffer only for background correction. Reporters were diluted in Reporter Stock Solution (PhenoCycler-Fusion Buffer, assay reagent, and 4 mM Hoechst 33342).

Before Flow Cell assembly, slides were transferred from storage buffer to PBS for 10 min. Flow Cells were assembled using the Akoya assembly device, adhered to dried slides, and equilibrated in PhenoCycler-Fusion Buffer for 10 min. Assembled Flow Cells were loaded into the PhenoCycler-Fusion Slide Carrier to interface with the microfluidics system.

Runs were configured in the PhenoCycler Experiment Designer with the appropriate number of cycles and optimized exposure times. Imaging was performed on the PhenoImager Fusion (v1.6.1) at 20× with automated focusing and tissue detection. Raw 16-bit images were processed using Akoya’s updated software that retains full 16-bit depth in the compiled qPTIFF files.

### Post-CODEX H&E Staining

After the CODEX run was finished, the Flow Cell was manually removed from the slide. The slide was then washed in PBS three times for 5 min and given to the SCCC Tissue Management Core for H&E staining and subsequent brightfield image acquisition at 40x magnification.

### Image Registration

The first step in SPACEMAP is image registration. H&E-stained images are registered to CODEX multiplex images to enable integrated analysis, as exemplified with datasets acquired at different magnifications (e.g., H&E at 40×, 0.2495 μm/pixel; CODEX at 20×, 0.5083 μm/pixel). The pipeline first preprocesses H&E images by resizing them to match CODEX resolution via area interpolation if needed, ensuring alignment compatibility across varying acquisition scenarios. The DAPI channel is then extracted from the CODEX image, handling diverse formats such as (channels, height, width) or (height, width, channels), using tifffile to serve as a structural reference. Affine transformation is computed with the palom library (version 2024.12.1), employing thumbnail-based coarse registration (level 2) with ∼10,000 keypoints followed by full-resolution refinement (level 0) and shift constraints for robust handling of large, crowded tissues. Registered H&E channels (converted to RGB if grayscale) are merged with all CODEX markers into pyramidal OME-TIFF files via dask (24) for memory-efficient, channel-wise processing, padding dimensions as required and preserving metadata like channel names, colors, and physical pixel sizes. This modular capability supports batch processing of multi-patient slides, artifact handling, and seamless integration with tools like QuPath, facilitating scalable spatial biology workflows.

### Separation of Regions of Interest (ROIs)

SPACEMAP optionally extracts ROIs from registered CODEX-H&E OME-TIFF files to support focused, region-specific analyses, particularly beneficial for slides containing multiple ROIs such as tissue microarrays (TMAs) with cores from various patients, multiple tissue types, or diverse samples per slide. Leveraging GeoJSON boundary annotations exported from QuPath, the pipeline parses bounding boxes matched by base name, cropping each ROI independently while validating dimensions and excluding invalid or out-of-bounds regions. This separation into individual pyramidal OME-TIFF files enables faster processing of large datasets by allowing parallel or batch-efficient handling of smaller, targeted images rather than entire slides. Processing occurs channel-by-channel with tifffile to manage memory for high-resolution datasets (>10^6^ pixels), generating outputs (e.g., 3 pyramidal levels, 256×256 tiles, deflate compression) that retain full metadata, including channel information and pixel calibration (e.g., 0.5083 μm/pixel). Customizable parameters like pyramid depth and tile size adapt to diverse image sizes, while features such as metadata inheritance and fallback handling ensure robustness across scenarios like incomplete annotations. This capability facilitates detailed downstream tasks in SPACEMAP, such as cell segmentation and spatial zonation, by producing QuPath-compatible files optimized for visualization and analysis.

### Background Subtraction for OME-TIFF Images using Blank Cycles

To address residual background signals and imaging artifacts in Akoya CODEX-processed multiplex images, arising from non-specific antibody binding, autofluorescence, or optical aberrations, we incorporated a blank imaging cycle using non-specific fluorophores matched to the experimental spectral channels. This cycle captures channel-specific noise profiles, including non-specific adhesion, system background, detector noise, and residual autofluorescence. SPACEMAP performs pixel-wise background subtraction on registered OME-TIFF files via a streamlined algorithm: parsing OME-XML metadata for channel and image properties; user-defined mapping of blank to marker channels; subtraction using corrected_pixel = max(0, marker_pixel - blank_pixel) with multi-resolution pyramid generation; and optional quality control visualizations. Employing memory-efficient streaming, tiled compression, and full metadata preservation, the process yields 16-bit OME-TIFF outputs retaining original resolution, calibration, and channel ordering (excluding blanks), suitable for downstream segmentation and phenotyping.

### Segmentation and Quality Control

SPACEMAP integrates two deep learning-based segmentation models: Cellpose and DeepCell (Mesmer), allowing users to select the approach best suited to their datasets, such as tissue microarrays or whole-slide images with varying resolutions and marker panels. Both models process a two-channel input: the first channel uses a nuclear stain (e.g., DAPI) for initial cell detection, while the second is a composite membrane image created via maximum intensity projection of user-specified markers (e.g., Pan-Cytokeratin, E. cadherin, Vimentin, CD45, CD44, aSMA, CD45RO, CD4, CD8, CD3e, CD31, CD20, CD11c, CD69, CD11b, CD163, CD38 and HLA-A) to refine boundaries and reduce background artifacts. Optional tissue masking, derived from pathologist-guided GeoJSON annotations in QuPath, can be applied to restrict segmentation to regions of interest, excluding artifacts or non-tissue areas for improved accuracy and efficiency. Segmentation leverages GPU acceleration via PyTorch or TensorFlow for scalable processing of large datasets, with outputs including a feature table (CSV) containing morphological metrics (e.g., area, eccentricity, solidity) and mean marker intensities per cell/nucleus, integer-labeled masks (TIFF or NPY), and cell/nuclear boundaries in GeoJSON format for visualization and spatial analysis in tools like QuPath. A key innovation is the QuPath-compatible GeoJSON format, which connects segmentation outputs to co-registered H&E and CODEX images for visual assessment and manual quality control. To determine the impact of training, ten random regions were selected from each tissue type (epithelia, stroma, and tumor) and nuclei were manually counted as the reference standard. For each region, we calculated the ratio of segmented cell counts (default or trained model) to the manual nuclear count, with values closer to one indicating better agreement. Ratios from default and trained models were compared for statistical significance.

### Cellpose

SPACEMAP supports pre-trained Cellpose models (e.g., ‘cyto’) or custom-trained models via the Cellpose v3.0.9 GUI and library, enabling adaptation to specific tissues or staining patterns to minimize false positives (e.g., debris detection) and maximize true positive cell capture. For instance, in colorectal cancer datasets, a custom model trained on annotated ROIs improved segmentation by optimizing parameters such as flow threshold (0.8) and automatic diameter estimation (set to 0), with channels configured as [1, 3] for cytoplasm (membrane composite) and nucleus. The pipeline prepares RGB input stacks (membrane in red, zeros in green, DAPI in blue) and extracts features using scikit-image’s regionprops, ensuring compatibility with downstream phenotyping.

### DeepCell

DeepCell Mesmer in SPACEMAP utilizes pre-trained models or user-provided trained models for simultaneous nuclear and whole-cell segmentation, with configurable compartments ("nuclear", "whole-cell", or "both") and primary output mask selection (e.g., "cell"). Input normalization scales nuclear and membrane channels to [0, 1], and batch processing (default size 4) handles large images efficiently. For example, in multiplexed tissue datasets, the "both" mode extracts separate features for cells and nuclei, storing them in distinct CSVs and GeoJSON files, facilitating analyses like nuclear morphology in immune cells.

### Annotations

SPACEMAP leverages the co-registered H&E and CODEX images to enable precise tissue annotation in QuPath, combining H&E’s superior morphological detail for structural context (e.g., tumor boundaries, stroma) with CODEX’s high-plex marker data for molecular phenotyping, resulting in more accurate compartment delineation than either modality alone (25). This registration allows pixel-level precision in annotations, overcoming limitations of pre-registration H&E-only annotation by pathologists, where tumor regions often include mixed stromal tissues or imprecise boundaries; post-registration, QuPath’s classifier integrates marker intensities and H&E morphologies for refined, accurate boundaries at the cellular level. Annotations are processed in a multi-stage workflow: initial classification, artifact removal and merging, zone generation, and cell-level spatial feature extraction, with outputs in QuPath-compatible GeoJSON format for seamless integration and manual refinement.

### Tissue Classification

Tissue compartments (e.g., epithelium, stroma, tumor, endothelia) are classified using QuPath’s Random Trees pixel classifier on the unified OME-TIFF images. Pre-processing applies Gaussian and Laplacian-of-Gaussian filters to enhance features, training the classifier on marker expression and morphology. Outputs are exported as GeoJSON annotations for downstream processing.

### Artifact Removal and Tissue Merging

Histological artifacts (e.g., folds, clots, debris) are manually annotated in QuPath and categorized (e.g., "artifact", "exclude", "blood_clot", etc) for exclusion. An automated Python script, utilizing Shapely for geometric operations (26) loads primary annotations, integrates endothelial regions, subtracts exclusion zones, and merges cleaned components into a unified "tissue" polygon, filtering fragments below 300,000 pixels² to eliminate less usable dissociated tissues. The novel output, a single QuPath-compatible GeoJSON file with cleaned subcomponents, facilitates efficient manual refinement, enhancing reproducibility over manual-only workflows.

### Tissue Zone Classification

Post-refinement, a second script crops high-resolution subcomponents to the master "tissue" polygon, ensuring alignment with quality-controlled boundaries. Spatial zones (e.g., tumor-stroma margins) are programmatically generated via buffering (e.g., 50 μm) and intersection, with flexibility to define multiple concentric zones inside or outside user-selected structures such as tumors or normal epithelia; for instance, zones can be created within the tumor (e.g., core, mid-layer, periphery) or in adjacent stroma (e.g., immediate invasive front, intermediate, distant), resulting in contiguous zones that touch at interfaces to enable detailed spatial analysis of cellular interactions across boundaries. Small fragments below 3,000 pixels² are filtered to promote tissue continuity by removing microstructural gaps or islands (e.g., stroma-like artifacts within epithelia or vice versa), minimizing classification artifacts and ensuring robust, cohesive compartments for downstream analysis. This automated zone creation, novel in its integration with co-registered modalities, produces final GeoJSON files for cell annotation.

### Spatial Feature Extraction

Segmented cells from Step 4 notebook are annotated by mapping centroids to GeoJSON compartments and zones using Shapely and SciPy’s cKDTree for efficient nearest-neighbor queries. Each cell receives labels for primary compartment (e.g., "tumor", "stroma"), ROI, zone (e.g., "zone_a" for tumor core), and tumor proximity (e.g., distances via k-d tree). Outputs are enriched CSVs per slide, with geometric contours in GeoJSON, enabling quantitative spatial analyses like immune infiltration at tumor margins.

### Batch Correction and Quality Control (QC)

SPACEMAP implements batch correction to mitigate technical variations across experimental runs while preserving biological signals, adapted into Python from the cyCombine framework (5). The pipeline processes annotated cell data from Step 4, incorporating optional metadata (e.g., treatment, timepoints) for downstream analysis. Key steps include pre-correction QC with histogram analysis for marker distributions and outlier detection via percentile thresholds (e.g., 99.99th for high expression, 0.01th for low); per-batch Z-score standardization using scikit-learn’s StandardScaler; SOM-based clustering (e.g., 8×8 grid, 100,000 iterations) with MiniSom to group similar cells across batches; per-cluster parametric ComBat correction using pyComBat (27) to adjust for batch effects; and value capping to constrain corrected expressions to original ranges, preventing overcorrection. Post-correction QC generates before/after visualizations (e.g., density plots, PCA, UMAP) to assess efficacy, with options for filtering overcorrected cells (e.g., >99th percentile correction distance) or UMAP outliers. Outputs include combined and individual corrected CSVs with raw and adjusted markers, QC reports, and processing summaries, enabling robust downstream phenotyping by ensuring comparable data across batches.

### Threshold Detection

Prior to phenotyping, marker-specific thresholds are automatically detected per batch or ROI using a custom QuPath script adapted from open-source extensions (28) (version 25.02.1), incorporating ImageJ’s auto-threshold algorithms (e.g., Triangle) (29) on DAPI or marker channels. The script processes annotations or full images at user-defined downsampling (e.g., 5×), with options for dark background correction and minimum thresholds, computing values via histogram analysis. A key advantage is the direct assignment of thresholds to OME-TIFF metadata and export to CSVs, enabling seamless integration with RESOLVE, SCIMAP, or other tools for reproducible analysis across modalities.

### Phenotyping

SPACEMAP integrates four phenotyping methods, RESOLVE (novel thresholding-based), Leiden clustering, Self-Organizing Maps (SOMs), and SCIMAP, within ensemble strategies to enhance robustness and reduce method-specific biases. Using annotated data from Step 4 (with batch-corrected markers for Leiden and SOM, and threshold-based processing for RESOLVE and SCIMAP), phenotyping assigns cell identities based on marker expression, with outputs including phenotype labels, confidence scores, and visualizations for validation. The ensemble approaches include a manual-reference model, where methods are compared to expert annotations and weighted by F1 scores in a machine learning classifier, and a reference-free consensus model, identifying high-confidence cells (agreement across ≥3 methods) for training a predictive model, both yielding superior F1 scores over individual methods.

### RESOLVE

RESOLVE, a novel thresholding-based phenotyping algorithm developed internally for this study, begins by subtracting batch- or ROI-specific thresholds from marker intensities to eliminate background noise (Supplementary Fig. 7). This is followed by optional flexible normalization steps, adapted from Hickey et al. (11), including marker-wise Z-scoring within groups (e.g., per batch), cell-wise Z-scoring, cumulative distribution function (CDF) transformation, and negative log scaling, to standardize expression values across datasets. Binary positivity flags are then assigned for each marker (positive if above threshold). Initial phenotypes are determined using logical combinations of these flags, such as OR for inclusive markers and for required co-expression (e.g., a regulatory T cell is classified as CD45+ AND CD3e+ AND CD4+ AND FoxP3+). In cases of conflicting assignments, a predefined hierarchy prunes broader categories in favor of more specific subtypes (e.g., prioritizing "regulatory T cell" over "helper T cell" or daughter cell over parental), with remaining ties resolved by selecting the phenotype with the highest normalized expression on its defining marker. This unsupervised thresholding combined with hierarchical logic provides a reproducible, bias-minimizing framework for accurate cell identification in multiplexed imaging data.

### Leiden Clustering

Automated cell phenotyping was performed using Leiden clustering, an unsupervised, graph-based workflow implemented in Python with the Scanpy library. The analysis began with batch-corrected protein marker expression data. This data was processed for dimensionality reduction, first using Principal Component Analysis (PCA). A k-nearest neighbor graph was constructed from these components, which then served as the input for both Uniform Manifold Approximation and Projection (UMAP) embedding and subsequent clustering. Cell populations were identified by applying the Leiden algorithm to this graph. The resulting clusters were annotated with biological cell phenotypes based on the mean expression of canonical markers, a process facilitated by UMAP visualizations and cluster expression heatmaps. All generated data, including cell phenotype labels and summary plots, were saved for downstream analysis.

### Self-Organizing Map (SOM)

Cell phenotyping was performed using a two-tiered clustering approach centered on Self-Organizing Maps, implemented in Python using the MiniSom and scikit-learn libraries. Initially, standardized expression data from selected phenotyping-protein markers were used to train a 15×15 SOM over 100 iterations partitioning cells into 225 micro-clusters. To create a more tractable set for annotation, the mean marker expression profiles of these 225 SOM nodes were then subjected to hierarchical agglomerative clustering using Ward’s linkage method with a Euclidean distance metric, grouping them into 100 final meta-clusters. Biological phenotypes were assigned to these meta-clusters through manual inspection of a clustered heatmap that visualized the Z-scored mean expression of canonical markers.

### SCIMAP

Cell phenotyping was conducted using a sequential gating workflow inspired by the SCIMAP toolkit and implemented in Python. Raw marker expression data for selected phenotyping-proteins were first log-transformed (log(x+1)). Each marker was then independently rescaled to a range of [0, 1] to normalize signals across the dataset, with scaling designed to incorporate automatically detected thresholds (from the earlier threshold detection method) such that values from 0 to 0.5 represent scaled log-transformed intensities below the threshold (background/negative expression), while values from 0.5 to 1 represent scaled log-transformed intensities above the threshold (positive expression). Following this normalization, cell identities were assigned by applying a pre-defined hierarchical gating strategy to the rescaled data. A cell was classified as positive for a given marker if its rescaled expression exceeded a gate of 0.5, and phenotypes were defined by unique combinations of positive and negative markers as specified in a predefined strategy file.

### Manual Reference Annotation

To establish the manual reference for model training and validation, two distinct sets of manual reference annotations were created with expert pathologist supervision using QuPath (MW and PG). First, to train the manual reference-weighted ensemble model (MR-ML), approximately 6,000 cells were randomly selected across multiple patient samples and tissue regions. For this process, the GeoJSON cell segmentation boundaries were imported into QuPath and overlaid on the multiplex immunofluorescence images. An expert pathologist then assigned a phenotype to each randomly selected cell based on the visual expression of canonical protein markers. This random sampling strategy ensured the creation of a diverse and representative dataset for training a robust classifier.

Second, a separate manual reference set was created for the specific purpose of validating the agreement between the MR-ML and Consensus-ML models and for generating high-resolution visualization figures. For this set, cells within a single, contiguous region of interest were annotated. This approach provided a spatially coherent ground truth for direct model comparison and visualization.

### Manual Reference Machine-Learning (MR-ML)

To enhance phenotyping accuracy and RESOLVE discordance among individual methods, we developed an ensemble machine learning model trained on a manually annotated reference dataset. A subset of random cells, around 6000, from multiple patients was first manually classified to serve as the ground truth. This reference set was then used to evaluate the performance of each of the four automated phenotyping methods (RESOLVE, SCIMAP, Leiden, and SOMs) by calculating their phenotype-specific F1-scores.

A Random Forest classifier was trained on the batch-corrected marker expression data from the manually annotated cells. To predict the phenotype of unannotated cells, the model integrated two components: (1) the direct prediction from the trained Random Forest classifier, and (2) a weighted vote from each of the four automated methods. The weights for this vote were derived directly from the phenotype-specific F1-scores, giving more influence on methods that performed better for a given cell type. The final phenotype assignment was determined by the class with the highest combined score from this weighted ensemble. This approach leverages the accuracy of a manual reference while systematically integrating the outputs of multiple algorithms to produce a robust and high-confidence final phenotype for every cell in the dataset.

### Consensus Machine-Learning (Consensus-ML)

As a rapid and robust alternative to manual annotation, we developed a reference-free ensemble phenotyping model that leverages consensus agreement among multiple automated methods. This approach identifies high-confidence training labels by selecting "consensus” cells for which at least three of the four primary phenotyping methods (RESOLVE, SCIMAP, Leiden, and SOMs) independently assigned the same phenotype label. This strategy generated a large and diverse training set (over 50% of cells in our dataset were consensus) without the time-intensive process of manual expert annotation.

A Random Forest classifier was then trained on the batch-corrected marker expression data using these consensus cells as ground truth. The trained model was subsequently applied to predict phenotypes for all non-consensus cells. Each prediction was assigned a confidence score based on the maximum class probability from the classifier, while consensus cells were assigned a confidence score of 1.0. This automated, consensus-driven approach provides a scalable and efficient method for achieving high-confidence, full-dataset phenotyping, yielding results comparable to the manual reference model but with significantly reduced turnaround time.

### Machine Learning Phenotyping Quality Control Assessment

To evaluate phenotyping performance, we assessed five complementary metrics: F1 score, which balances precision and recall; accuracy, mearing the proportion of correctly classified cells; recall, reflecting sensitivity to true positive assignments; precision, indicating specificity of predictions and Cohen’s Kappa, quantifying agreement beyond chance **(Fig S6 and Fig S7)**.

To assess the reliability of automated phenotyping methods, we calculated Cohen’s kappa (κ) to measure agreement between each computational method and manual expert annotations, accounting for chance agreement and class distribution imbalances. Cohen’s kappa values of 0.6-0.8 indicate substantial agreement, while values >0.8 represent almost perfect agreement between methods and ground truth annotations.

### Statistics

All statistical analyses were performed using GraphPad Prism (GraphPad Software, San Diego, CA). Differences between two related (paired) groups were analyzed using unpaired t-tests. Significance was determined at a p<0.05.

## Acknowledgments

This work was supported by the Damon Runyon Cancer Research Foundation, Apexigen Inc, CPRIT Recruitment of First-Time, Tenure-Track Faculty Members RR170051, the University of Texas Southwestern Medical Center (UTSW) Disease-Oriented Scholars Program, UTSW Harold C.Simmons Comprehensive Cancer Center Translational Pilot Award, and National Cancer Institute 1R01CA283953 (TAA), and Mary Kay Ash Foundation (SD). We acknowledge the Harold C. Simmons Comprehensive Cancer Center (SCCC) Tissue Management Shared Resource and the UTSW Radiation Oncology Clinical Research Office. We thank the patients who trusted the team to enroll in the study and shared tissue samples for this study.

## References

1. Tan WCC, Nerurkar SN, Cai HY, Ng HHM, Wu D, Wee YTF, et al. Overview of multiplex immunohistochemistry/immunofluorescence techniques in the era of cancer immunotherapy. Cancer Commun. 2020 Apr;40(4):135–53.

2. Catching up with multiplexed tissue imaging. Nat Methods. 2022 Mar;19(3):259–259.

3. Sheng W, Zhang C, Mohiuddin TM, Al-Rawe M, Zeppernick F, Falcone FH, et al. Multiplex Immunofluorescence: A Powerful Tool in Cancer Immunotherapy. Int J Mol Sci. 2023 Feb 4;24(4):3086.

4. Bollhagen A, Bodenmiller B. Highly Multiplexed Tissue Imaging in Precision Oncology and Translational Cancer Research. Cancer Discov. 2024 Nov 1;14(11):2071–88.

5. Pedersen CB, Dam SH, Barnkob MB, Leipold MD, Purroy N, Rassenti LZ, et al. cyCombine allows for robust integration of single-cell cytometry datasets within and across technologies. Nat Commun. 2022 Mar 31;13(1):1698.

6. Schürch CM, Bhate SS, Barlow GL, Phillips DJ, Noti L, Zlobec I, et al. Coordinated Cellular Neighborhoods Orchestrate Antitumoral Immunity at the Colorectal Cancer Invasive Front. Cell. 2020 Sept;182(5):1341–1359.e19.

7. Wilson CM, Ospina OE, Townsend MK, Nguyen J, Moran Segura C, Schildkraut JM, et al. Challenges and Opportunities in the Statistical Analysis of Multiplex Immunofluorescence Data. Cancers. 2021 June 17;13(12):3031.

8. Ma J, Xie R, Ayyadhury S, Ge C, Gupta A, Gupta R, et al. The multimodality cell segmentation challenge: toward universal solutions. Nat Methods. 2024 June;21(6):1103–13.

9. Bai Y, Zhu B, Rovira-Clave X, Chen H, Markovic M, Chan CN, et al. Adjacent Cell Marker Lateral Spillover Compensation and Reinforcement for Multiplexed Images. Front Immunol. 2021 July 5;12:652631.

10. Rojas F, Hernandez S, Lazcano R, Laberiano-Fernandez C, Parra ER. Multiplex Immunofluorescence and the Digital Image Analysis Workflow for Evaluation of the Tumor Immune Environment in Translational Research. Front Oncol. 2022 June 27;12:889886.

11. Hickey JW, Tan Y, Nolan GP, Goltsev Y. Strategies for Accurate Cell Type Identification in CODEX Multiplexed Imaging Data. Front Immunol. 2021 Aug 13;12:727626.

12. Tan Y, Kempchen TN, Becker M, Haist M, Feyaerts D, Xiao Y, et al. SPACEc: A Streamlined, Interactive Python Workflow for Multiplexed Image Processing and Analysis [Internet]. Bioinformatics; 2024 [cited 2025 Sept 24]. Available from: http://biorxiv.org/lookup/doi/10.1101/2024.06.29.601349

13. Kuswanto W, Nolan G, Lu G. Highly multiplexed spatial profiling with CODEX: bioinformatic analysis and application in human disease. Semin Immunopathol. 2023 Jan;45(1):145–57.

14. Bruhns M, Schleicher JT, Wirth M, Zago M, Babaei S, Claassen M. Effects of segmentation errors on downstream-analysis in highly-multiplexed tissue imaging. Lorenzo G, editor. PLOS Comput Biol. 2025 Sept 15;21(9):e1013350.

15. Van Valen DA, Kudo T, Lane KM, Macklin DN, Quach NT, DeFelice MM, et al. Deep Learning Automates the Quantitative Analysis of Individual Cells in Live-Cell Imaging Experiments. Meier-Schellersheim M, editor. PLOS Comput Biol. 2016 Nov 4;12(11):e1005177.

16. Stringer C, Wang T, Michaelos M, Pachitariu M. Cellpose: a generalist algorithm for cellular segmentation. Nat Methods. 2021 Jan;18(1):100–6.

17. Vierdag WMAM, Saka SK. A perspective on FAIR quality control in multiplexed imaging data processing. Front Bioinforma. 2024 Feb 9;4:1336257.

18. Kang Z, Szabo A, Farago T, Perez-Villatoro F, Junquera A, Shah S, et al. Tribus: semi-automated discovery of cell identities and phenotypes from multiplexed imaging and proteomic data. Gao X, editor. Bioinformatics. 2025 Mar 4;41(3):btaf082.

19. Traag VA, Waltman L, Van Eck NJ. From Louvain to Leiden: guaranteeing well-connected communities. Sci Rep. 2019 Mar 26;9(1):5233.

20. Kohonen T. Self-organized formation of topologically correct feature maps. Biol Cybern. 1982;43(1):59–69.

21. Nirmal AJ, Sorger PK. SCIMAP: A Python Toolkit for Integrated SpatialAnalysis of Multiplexed Imaging Data. J Open Source Softw. 2024 May 29;9(97):6604.

22. Tang J, Du W, Shu Z, Cao Z. A generative benchmark for evaluating the performance of fluorescent cell image segmentation. Synth Syst Biotechnol. 2024 Dec;9(4):627–37.

23. Radtke AJ, Postovalova E, Varlamova A, Bagaev A, Sorokina M, Kudryashova O, et al. Multi-omic profiling of follicular lymphoma reveals changes in tissue architecture and enhanced stromal remodeling in high-risk patients. Cancer Cell. 2024 Mar;42(3):444–463.e10.

