## Supplementary Figures for "Resolving phenotyping discordance with SPACEMAP, an integrated machine learning framework"

**Supplementary Materials**

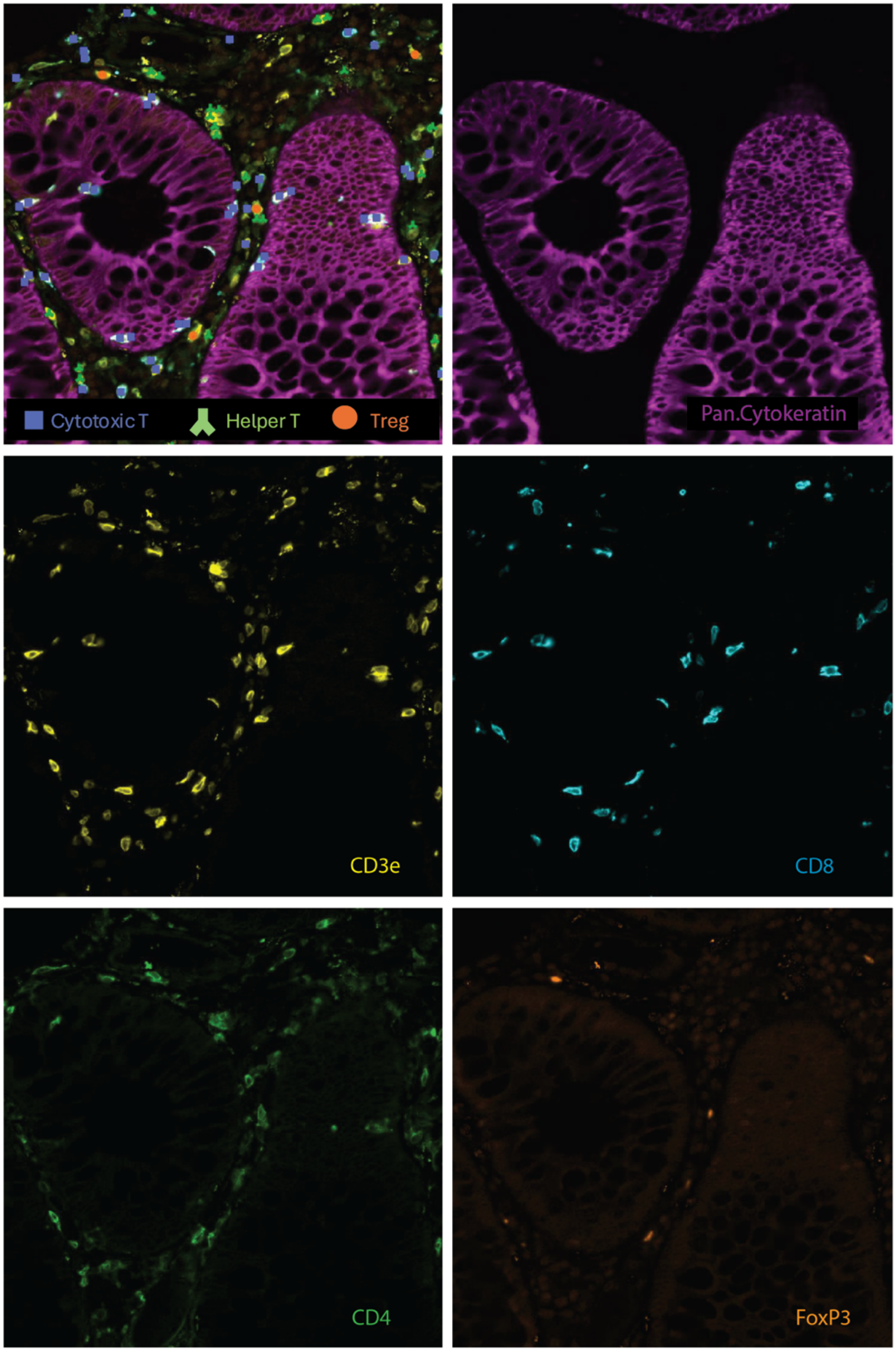


**Figure S1. RESOLVE Reveals Immune Cells in Epithelium.** Representative RESOLVE phenotyping results showing immune cell populations localized within epithelial regions. Panels display markers for pan-cytokeratin, CD3e, CD8, CD4, and FoxP3, illustrating the presence of cytotoxic T cells, helper T cells, and regulatory T cells within the epithelial compartment.

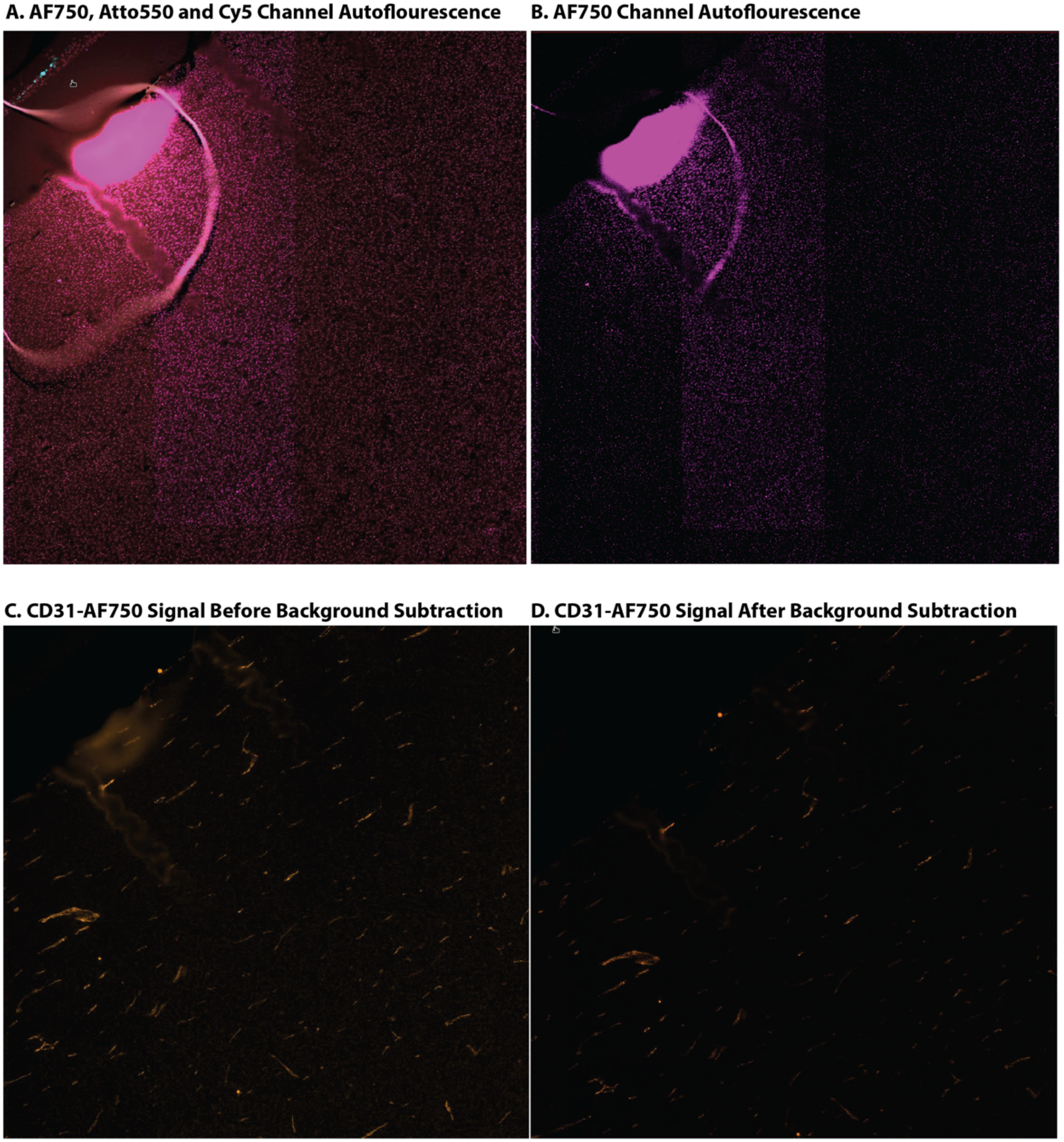


**Figure S2. Background Subtraction Improves Signal Detection. a** Representative image of a blank cycle acquisition cycle showing background autofluorescence in Alexa Fluor 750 (AF750), Atto550 and Cy5 channels during CODEX imaging. **b** Representative image showing backgeound autofluorescence in the AF750 channel during CODEX acquisition. **c** CD31 conjuagted to AF750 showing signal prior to background subtraction, highlighting elevated nonspecific fluorescence. **d** CD31conjuagted to AF750 showing signal following background subtraction, demonstrating improved contrast and specific marker visualization.

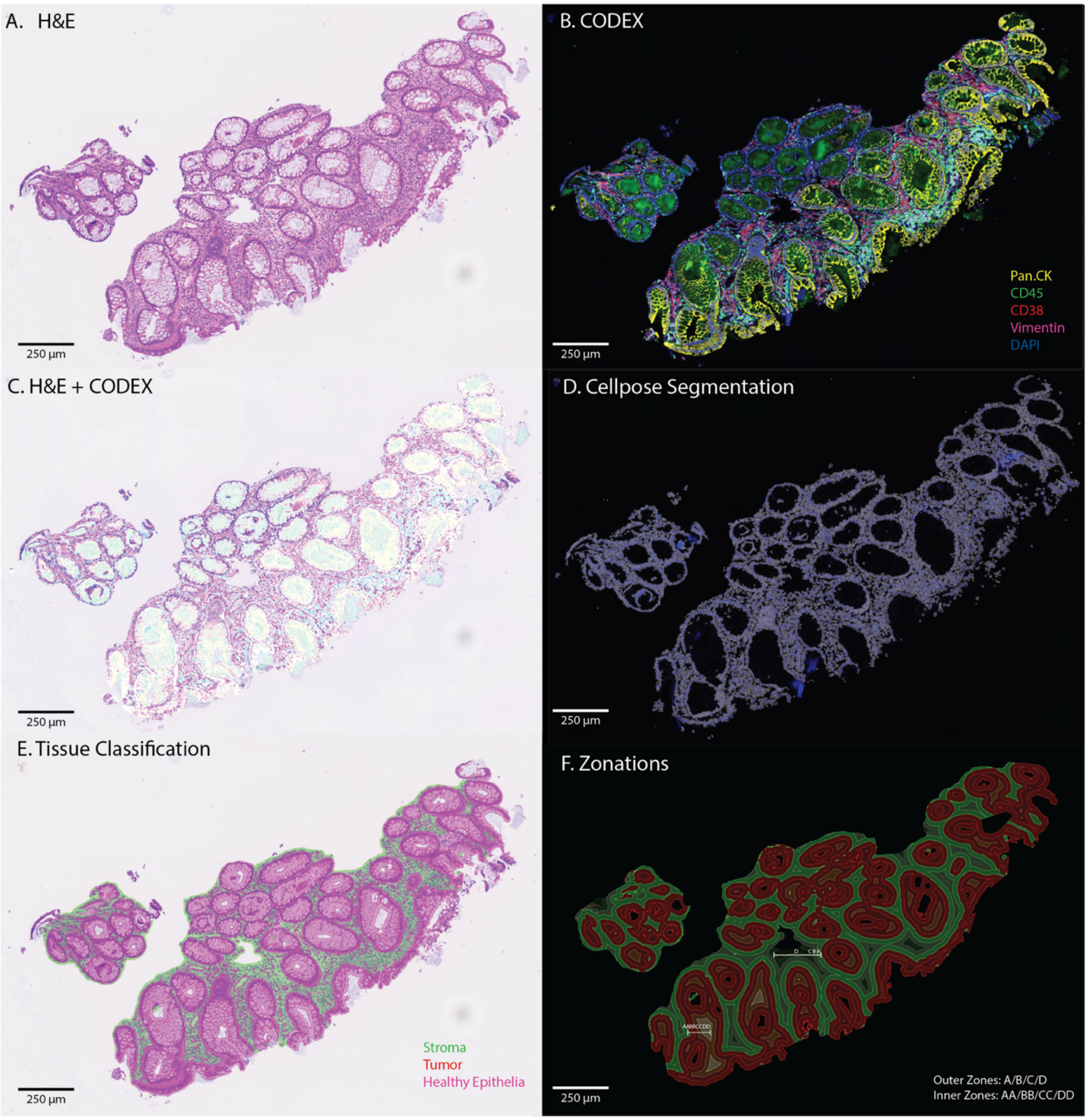


**Figure S3. Image Co-Registration of Healthy Tissue. a** The original H&E image, **b** Fluorescent channels overlay: Pan.Cytokeratin, α-Smooth Muscle Actin, CD45, Vimentin and DAPI **c** the final, co-registered OME-TIFF with the H&E image overlaid as a transparent layer. This visually demonstrates the accuracy of the alignment. **d** Trained cellpose segmentation overlaid on the DAPI channel **e** Tissue classification distinguishing tumor regions, stroma and healthy tissue **f** Zonation map indicating inner (AA, BB, CC, DD) and outer (A, B, C, D) regions across the tissue.

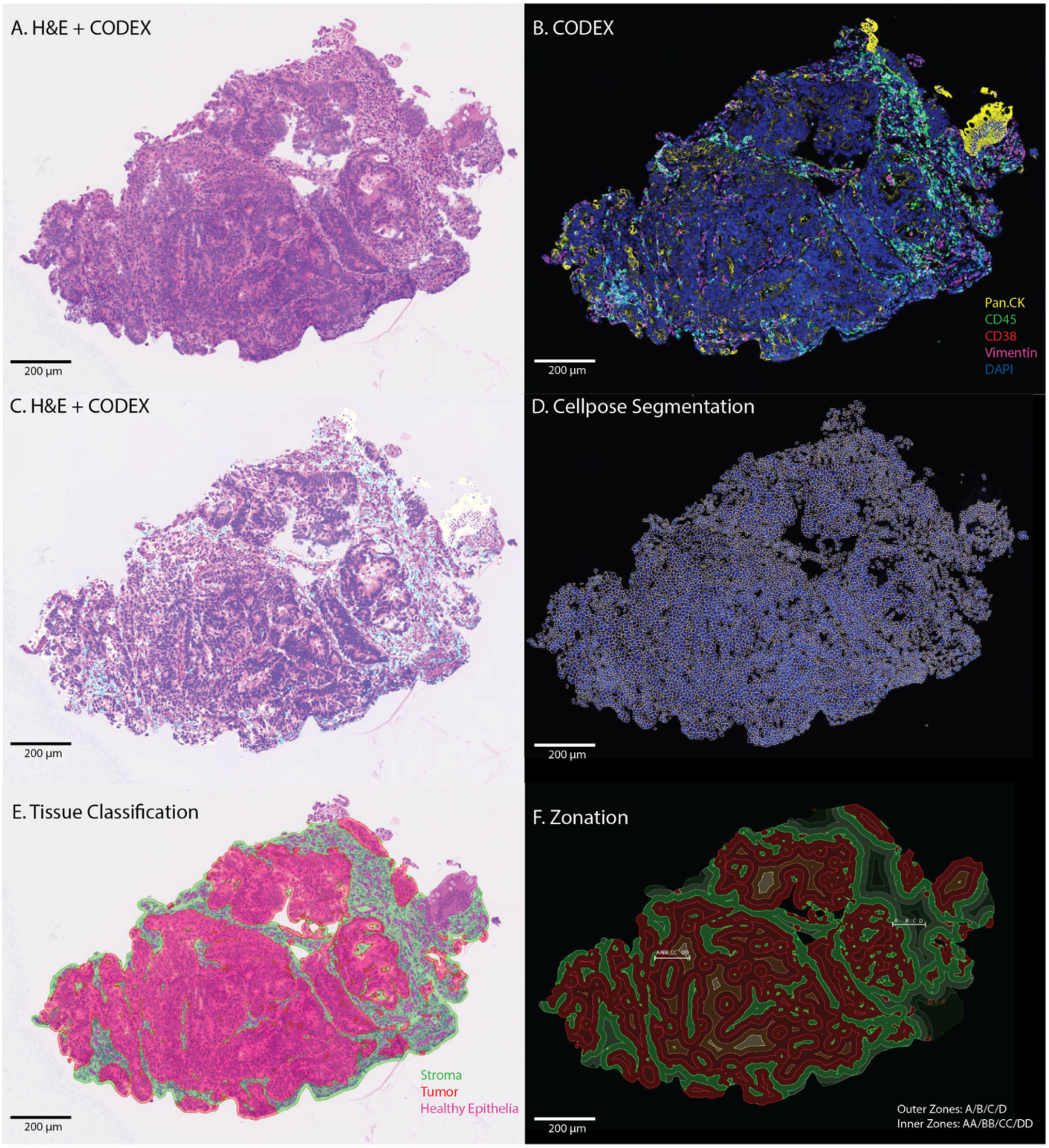


**Figure S4. Image Co-Registration of Tumor Tissue. a** The original H&E image, **b** Fluorescent channels overlay: Pan.Cytokeratin, α-Smooth Muscle Actin, CD45, Vimentin and DAPI **c** the final, co-registered OME-TIFF with the H&E image overlaid as a transparent layer. This visually demonstrates the accuracy of the alignment. **d** Trained cellpose segmentation overlaid on the DAPI channel **e** Tissue classification distinguishing tumor regions, stroma and healthy tissue **f** Zonation map indicating inner (AA, BB, CC, DD) and outer (A, B, C, D) regions across the tissue.

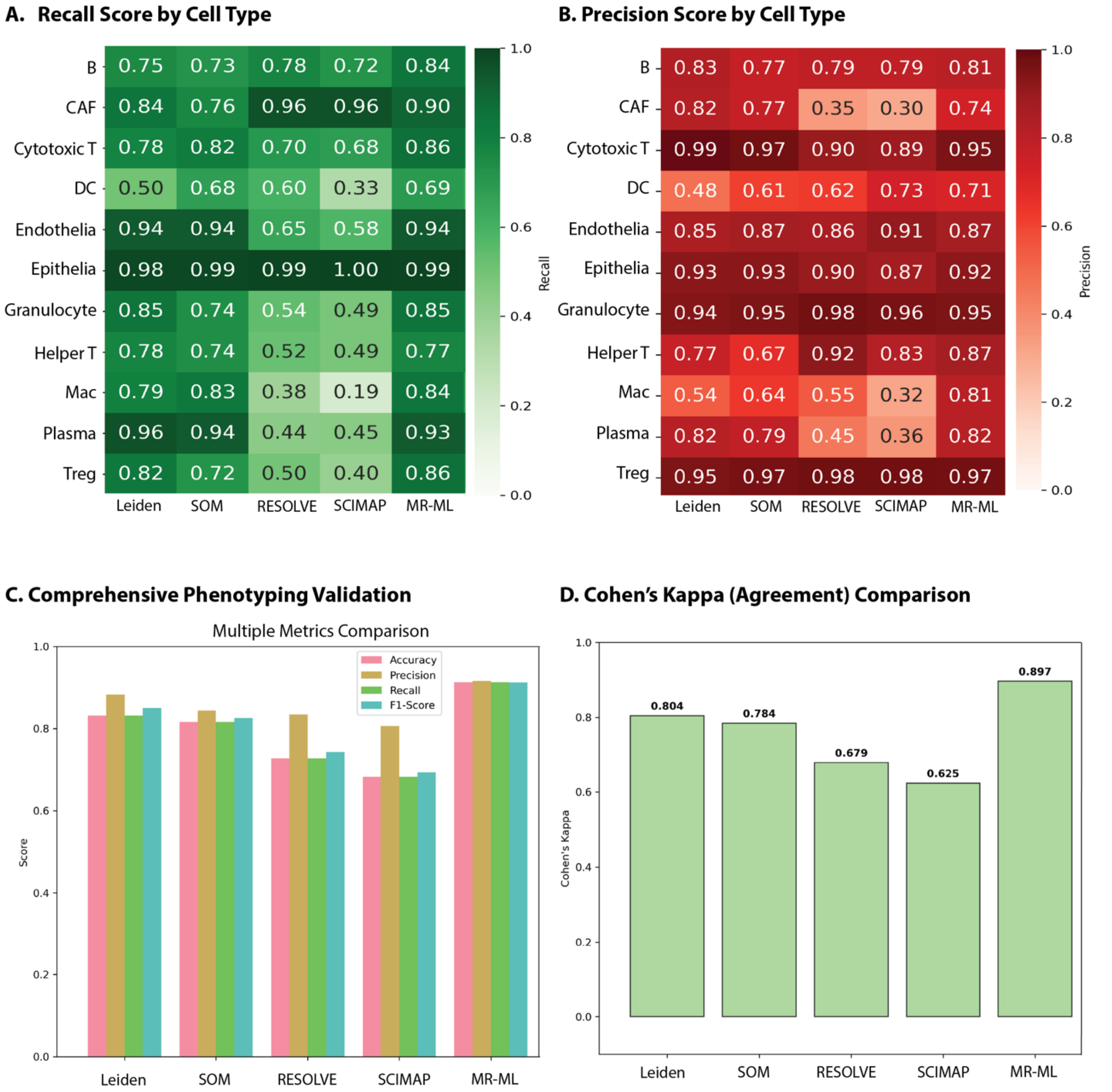


**Figure S5. Comparative Evaluation of Phenotyping Performance Across Methods.**
Performance metrics for Leiden, SOM, RESOLVE, and SCIMAP phenotyping approaches are shown, along with comparison to the weighted manual reference–machine learning approach. **a** Recall scores by cell type across all methods. **b** Precision by cell type across all methods. **c** Bar graphs comparing overall accuracy, precision, recall, and F1-score across the five methods. **d** Cohen's kappa analysis showing agreement levels among methods relative to the manual reference–machine learning standard.

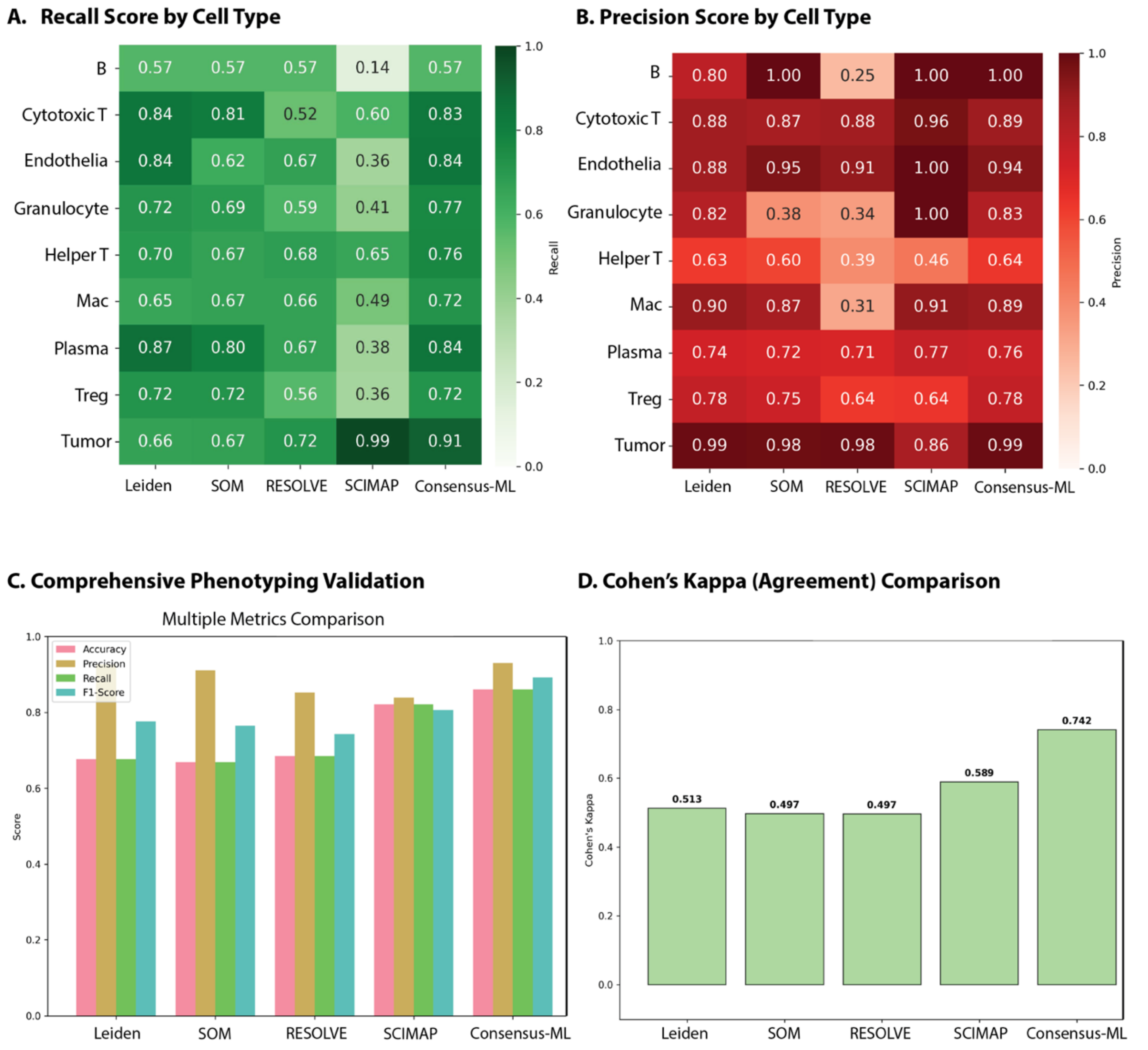


**Figure S6. Comparative Evaluation of Phenotyping Performance on the Nolan Dataset.** Performance metrics for Leiden, SOM, RESOLVE, and SCIMAP phenotyping approaches are shown, along with comparison to the weighted manual reference–machine learning approach. **a** Recall scores by cell type across all methods. **b** Precision by cell type across all methods. **c** Bar graphs comparing overall accuracy, precision, recall, and F1-score across the five methods. **d** Cohen's kappa analysis showing agreement levels among methods relative to the manual reference–machine learning standard.

**A. Channels Before and After Background Subtraction**

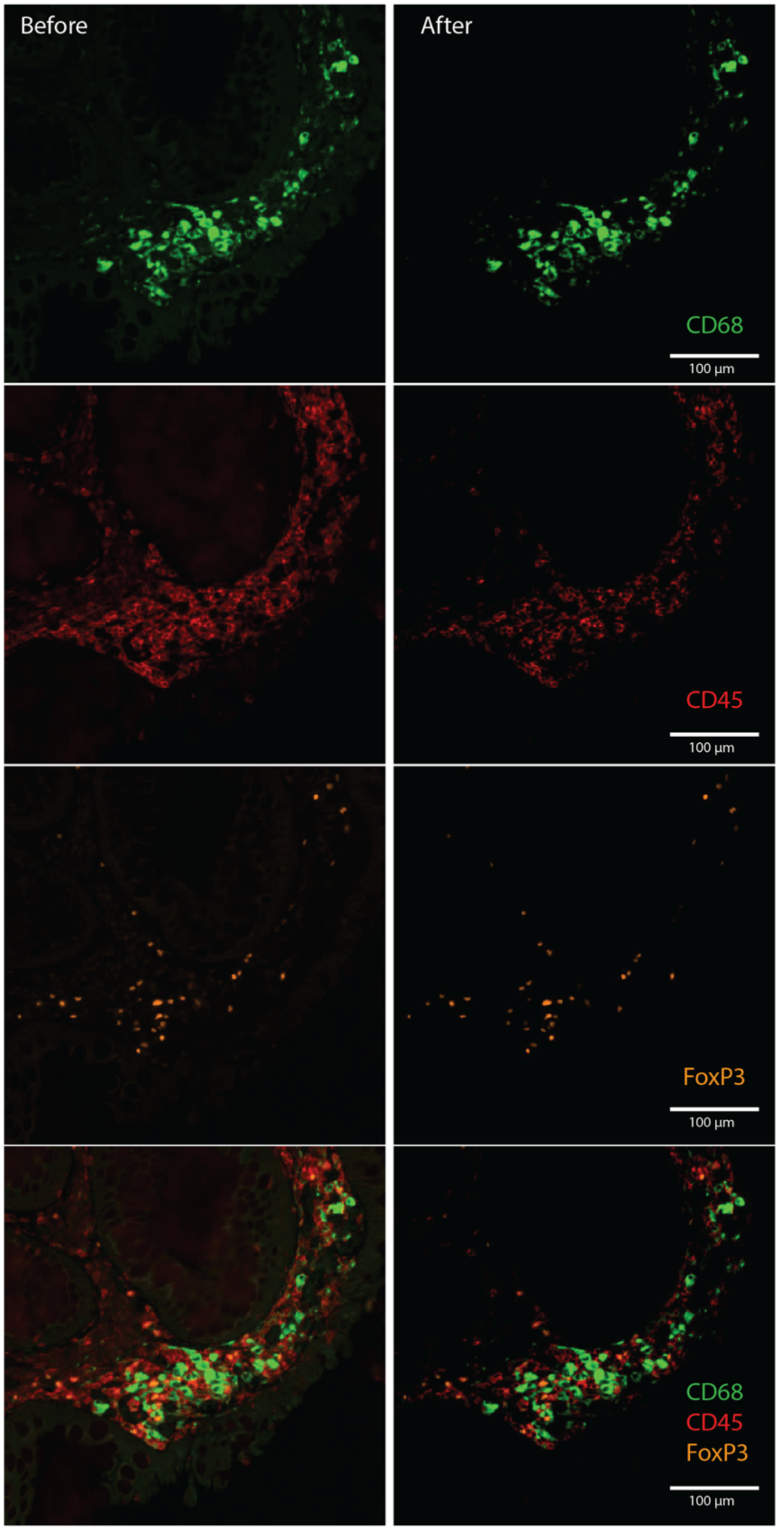


**Figure S7. Threshold-Based Background Subtraction for Phenotyping Accuracy.** Background subtraction, an optional preprocessing step during phenotyping in RESOLVE and SCIMAP, is illustrated for CD68, CD45, and FoxP3 channels. Shown are representative images before and after threshold-based background subtraction, highlighting improved signal clarity and reduced nonspecific fluorescence. The final panel displays an overlay of all channels, demonstrating enhanced visualization of distinct immune populations following background correction.

| Target | Clone | Vendor | Barcode | Fluorphore | Dilution | Exp. time (ms) | RRID |
|---|---|---|---|---|---|---|---|
| aSMA* | polyclonal | Abcam | BX041 | Atto550 | 1:200 | 100 | AB_2223021 |
| Axl* | C89E7 | CST | BX037 | Atto550 | 1:100 | 250 | AB_11217435 |
| CD1*c | L161 | Biolegend | BX002 | Atto550 | 1:100 | 250 | AB_1088996 |
| CD3e | EP449E | Akoya | BX045 | AF647 | 1:400 | 225 | AB_2915971 |
| CD4 | EPR6855 | Akoya | BX003 | Cy5 | 1:200 | 175 | AB_3094499 |
| CD8 | C8/144B | Akoya | BX026 | Atto550 | 1:200 | 200 | AB_2915960 |
| CD11b | EP1345Y | Abcam | BX010 | AF647 | 1:100 | 150 | Cat #ab187537 |
| CD11c* | D3V1E | CST | BX024 | Cy5 | 1:100 | 150 | AB_2943244 |
| CD15* | HI98 | Biolegend | BX029 | Atto550 | 1:3200 | 100 | AB_314194 |
| CD20 | L26 | Akoya | BX007 | AF750 | 1:800 | 200 | AB_2915939 |
| CD31 | EP3095 | Akoya | BX001 | AF750 | 1:200 | 200 | AB_2915935 |
| CD38 | E7Z8C | Akoya | BX089 | Atto550 | 1:4000 | 250 | AB_3082976 |
| CD40 | EPR20735 | Abcam | BX030 | AF647 | 1:200 | 250 | AB_2889383 |
| CD44 | IM7 | Akoya | BX005 | Atto550 | 1:600 | 150 | AB_2936081 |
| CD45* | 2D1 | Biolegend | BX046 | AF647 | 1:100 | 200 | AB_2566240 |
| CD45RO | UCHL1 | Akoya | BX017 | Atto550 | 1:600 | 200 | AB_2895053 |
| CD68 | KP1 | Akoya | BX015 | Cy5 | 1:1000 | 100 | AB_2935894 |
| CD69* | poly | R&D Systems | BX032 | Atto550 | 1:600 | 250 | AB_355231 |
| CD107a | H4A3 | Akoya | BX006 | Cy5 | 1:800 | 100 | AB_3474468 |
| CD141* | E7Y9P | CST | BX020 | Atto550 | 1:1600 | 150 | Cat No #34149 |
| CD163* | EDHu-1 | Novus | BX036 | Cy5 | 1:200 | 100 | AB_714951 |
| CD206* | polyclonal | R&D Systems | BX035 | Atto550 | 1:200 | 150 | AB_2063019 |
| E-cadherin | 4A2C7 | Akoya | BX014 | Atto550 | 1:400 | 150 | AB_2895057 |
| Fibronectin (FN1)* | F1 | Abcam | BX028 | Atto550 | 1:200 | 175 | AB_2895045 |
| FOXp3* | 236A/E7 | Thermo Fisher | BX027 | Cy5 | 1:100 | 100 | AB_467556 |
| gH2AX* | 2F3 | Biolegend | BX054 | AF647 | 1:200 | 100 | AB_315795 |
| HLA-A | EP1395Y | Akoya | BX004 | AF750 | 1:200 | 200 | AB_3678453 |
| HLA-DR | EPR3692 | Akoya | BX033 | AF647 | 1:100 | 150 | AB_3080864 |
| iNOS | SP126 | Akoya | BX023 | Atto550 | 1:100 | 250 | AB_3676531 |
| Ki67 | B56 | Akoya | BX047 | Atto550 | 1:400 | 150 | AB_2895046 |
| Pan-Cytokeratin (PanCK) | AE-1/AE-3 | Akoya | BX019 | AF750 | 1:200 | 100 | AB_3083456 |
| PCNA* | PC10 | Abcam | BX031 | AF647 | 1:3200 | 100 | AB_303394 |
| PDGFR*a | D13C6 | CST | BX021 | Cy5 | 1:100 | 150 | AB_10692773 |

| | | | | | | | |
|---|---|---|---|---|---|---|---|
| PD-L1 / CD274 | 73-10 | Akoya | BX043 | AF647 | 1:200 | 200 | Cat No #4550072 |
| Podoplanin | NC-08 | Akoya | BX0121 | Atto550 | 1:400 | 200 | AB_3082979 |
| S100A9* | D5O6O | CST | BX042 | Cy5 | 1:800 | 100 | AB_2799053 |
| Vimentin* | O91D3 | Biolegend | BX016 | AF647 | 1:3200 | 100 | AB_2565911 |

**Table 1. CODEX Antibody Panel.** A full list of all antibodies used in the CODEX panel, including the target, clone, vendor, Akoya-barcode, barcode-corresponding fluorophore, dilution factor, exposure time and identifier.

Akoya = Akoya Biosciences, CST = Cell Signaling Technology, Novus = Novus Biologicals.* = antibodies manually conjugated.
